# The Genetic Discovery (GeDi) resource: Deep characterization of a multi-ancestry panel of human reference iPSCs for functional genomic studies

**DOI:** 10.64898/2026.10.14.759173

**Authors:** Adrian Melo-Carrillo, Enrica Pellegrino, Parithi Balachandran, Mitch Kostich, Liz Werren, Cole Lorig, Angelina Kendra, Juliana Alcoforado Diniz, Katherine Merisotis, Jessica Garofalo, Stephen Grubb, Christine R. Beck, Paul Robson, William C. Skarnes

## Abstract

Human induced pluripotent stem cells (iPSCs) are essential tools for disease modelling, drug discovery, and functional genomics, yet the field remains anchored to a small number of reference lines, limiting our understanding of how genetic background shapes cellular phenotypes across human populations. Here we describe the Genetic Discovery iPSC panel (GeDi), eight clonal iPSC lines reprogrammed from male and female donors that broadly span global human population diversity. Ancestry analysis confirmed that each line’s genetic profile is concordant with donor self-reported ancestry. All GeDi lines are pluripotent and passed comprehensive quality control, demonstrating Sendai virus clearance, intact p53 pathway integrity, reproducible proliferative capacity, and compatibility with standardized genome editing workflows. Transcriptome profiling across five differentiated lineages covered 80% of annotated protein-coding genes and revealed stable donor-intrinsic variability in gene expression within each lineage. Haplotype-resolved genome assemblies were used to catalog structural variants and regions of absence of heterozygosity invisible to short-read sequencing. Whole-genome methylation analysis confirmed successful epigenetic reprogramming, catalogued the status of imprinted loci and captured the early dynamics of erosion of X chromosome inactivation (XCI) in female lines. The GeDi panel is available from The Jackson Laboratory as a quality-controlled resource designed to support functional genomic studies across varied genetic backgrounds.

## Introduction

Human induced pluripotent stem cells (iPSCs) have transformed biomedical research by providing an unlimited, patient-derived source of cells that can be differentiated into virtually any cell type of the human body (Takahashi, Tanabe et al. 2007, Yu, Vodyanik et al. 2007). Their capacity for indefinite self-renewal, combined with improved genome editing technologies, has made iPSCs essential for disease modeling, drug discovery, and functional genomics (Hockemeyer and Jaenisch 2016, Shi, Inoue et al. 2017). Despite the growing number of available lines, the field has converged on a small number of widely adopted reference lines — most notably KOLF2.1J, a male donor line of European ancestry used as a standard reference for large-scale genome editing initiatives (Pantazis, Yang et al. 2022), and WTC-11, a male donor line of East Asian ancestry valued for its high cloning efficiency and genomic stability (Miyaoka, Chan et al. 2014). While these lines have been invaluable for establishing reproducible workflows, their dominance means that the majority of functional genomic studies are conducted in the context of a narrow slice of human genetic diversity, limiting our ability to understand how genetic background shapes cellular phenotypes, gene regulation, and disease susceptibility across human populations (Popejoy and Fullerton 2016, Sirugo, Williams et al. 2019).

Individuals differ in their repertoire of common and rare variants, structural rearrangements, epigenetic landscapes, and downstream regulation of gene expression — differences that can profoundly influence cellular phenotypes, disease risk, and drug response (Genomes Project, Auton et al. 2015). This genomic diversity largely reflects the demographic and evolutionary histories of human populations, including migration, admixture, and adaptation to local environments (Campbell and Tishkoff 2008). African-ancestry populations harbor the greatest genetic diversity of any human group, a consequence of the longer history of human habitation on the continent and the serial founder effects that accompanied the out-of-Africa migration (Ramachandran, Deshpande et al. 2005, Campbell and Tishkoff 2008, Tishkoff, Reed et al. 2009), yet remain among the most underrepresented in iPSC studies. Initiatives such as the Human iPSC Initiative (HipSci; (Kilpinen, Goncalves et al. 2017), the iPSC Collection for Omic Research (iPSCORE; (Panopoulos, D’Antonio et al. 2017), and others have expanded the number of available lines, however these collections are still limited in genetic diversity, largely being derived from European and East Asian donors (Ghosh, Nehme et al. 2022). In addition, systematic characterization of iPSCs at the population level is generally lacking, and such characterization should span as much of global human diversity as possible to enable robust functional genomic use. Critically, the use of genetically diverse lines under identical experimental conditions also enables the separation of robust, lineage-defining biological signals from donor-specific variation — a distinction that is inaccessible when working with a single reference line.

Here we describe the first entries into the Genetic Discovery iPSC panel (GeDi), a collection of eight clonal iPSC lines reprogrammed from peripheral blood mononuclear cells (PBMCs) of healthy donors broadly spanning global human genetic diversity, comparing them to two well-characterized reference lines, KOLF2.1J and its edited derivative KOLF2.2J. All GeDi lines were subjected to a comprehensive quality control pipeline. Importantly, each of the eight lines were deeply characterized for genomic, transcriptomic, and epigenomic integrity and variation, with haplotype-resolved assemblies generated for each line. The GeDi panel and accompanying genomic data are maintained at The Jackson Laboratory as a community resource designed to support functional genomic studies across genetically distinct backgrounds.

## Results

### Reprogramming and selection of clonal sublines

Donor recruitment was designed, based on self-reported race and ethnicity, to maximize genetic heterogeneity while still having representation across major human populations (see methods for cohort recruitment details). PBMCs were isolated from whole blood of 23 donors by density gradient centrifugation and were reprogrammed by integration-free Sendai virus delivery of reprogramming factors (Chou, Mali et al. 2011) using a modified workflow (**Figure 1A**). Briefly, erythroid progenitor cells were expanded, transduced with CytoTune II virus, and cultured on a fibroblast feeder layer or feeder-free in ReproTeSR media. iPSC colonies were pooled and propagated feeder-free in TeSR Plus media for three passages, then adapted to StemFlex media for two passages prior to single-cell subcloning and the isolation of clonal iPSC lines. The rationale for this modified protocol was to gradually adapt reprogrammed cells to feeder-free conditions in StemFlex media, our preferred media for the culture, single-cell subcloning and genome editing of KOLF2-derived reference iPSC lines (Skarnes, Pellegrino et al. 2019). We were successful in generating iPSC pools from each of the 23 donor PBMC samples. iPSCs derived from female donors were designated GeDiX; lines from male donors were designated GeDiY; the 3-digit number refers to the donor from which the line is derived and is randomly generated in the de-identification process. We chose to initially subclone lines from 12 donor iPSC pools, selecting lines to balance sex and genetic diversity within the original 23-donor cohort, using self-reported race and ethnicity as an approximate but practical proxy for genetic ancestry.

**Figure 1.**
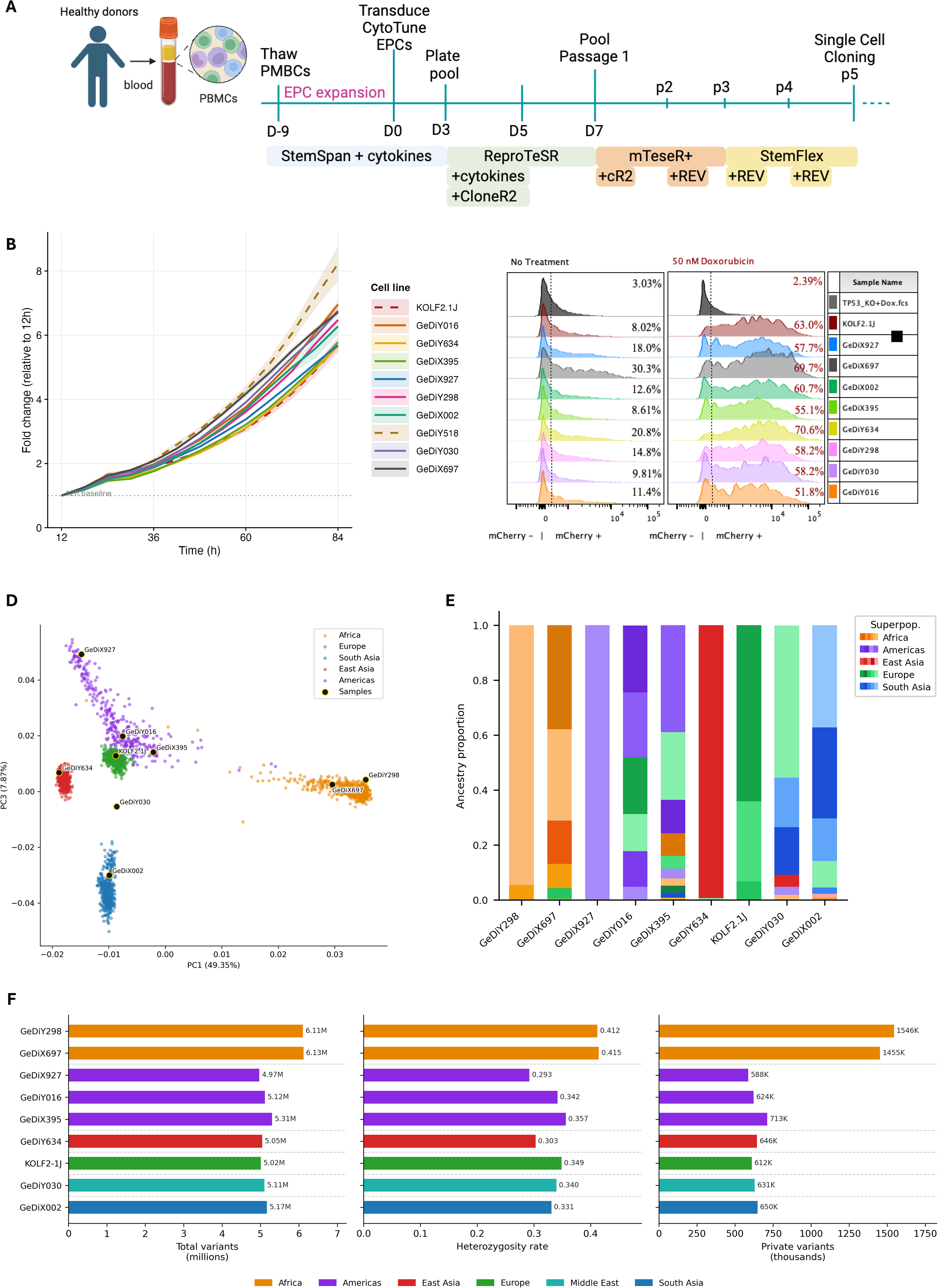
Characterization of GeDi iPSC lines. **(A)** Schematic overview of the GeDi iPSC reprogramming and quality control pipeline. PBMCs from healthy donors representing diverse population groups were reprogrammed using CytoTune Sendai virus. Following EPC expansion and transduction, cells were pooled and passage through defined media conditions before single-cell cloning at passage 5. **(B)** Growth curves of GeDi lines and KOLF2.1J measured by IncuCyte live-cell imaging over 80 hours (4X objective). Lines represent fold change in confluency relative to the 12 h timepoint. Shaded bands indicate 95% confidence intervals across three independent experiments. GeDiY518.1J (dashed orange) shows the fastest proliferation across the panel. **(C)** p53 DNA damage response assessed by mCherry reporter assay. Flow cytometry histograms show mCherry fluorescence under no treatment (left) and 50 nM doxorubicin treatment for 24 hours (right) for all GeDi lines, KOLF2.1J, and a TP53 KO control. Percentages indicate mCherry+ cells. All GeDi lines and KOLF2.1J show robust mCherry induction upon doxorubicin treatment, confirming intact p53 signaling. **(D)** Principal component analysis (PCA) of GeDi samples and KOLF2.1J projected onto the 1000 Genomes Project reference panel (PC1 versus PC3). Reference individuals are coloured by superpopulation (African, Americas, East Asian, European, and South Asian). GeDi samples are highlighted and labelled. GeDiY030 does not cluster with any reference superpopulation, consistent with Middle Eastern ancestry absent from the reference panel (Table S8). **(E)** Admixture plots for the eight GeDi individuals and KOLF2.1J. Each vertical bar represents a single individual; colors indicate inferred ancestry proportions across 26 reference populations from the 1000 Genomes Project (Table S3). Populations within each superpopulation are depicted as shades of the same base color. **(F)** Summary of genetic diversity metrics across GeDi samples and KOLF2.1J. Horizontal bar plots show total variants (millions), heterozygosity rate, and private variants (thousands) per individual. Samples are grouped and color-coded by superpopulation (African, Americas, East Asian, European, Middle Eastern, and South Asian). African samples exhibit the highest genetic diversity across all three metrics, consistent with established patterns of human genetic variation.

Within a pool of reprogrammed iPSC cells, we expect to find germline (Sudmant, Rausch et al. 2015, Logsdon, Ebert et al. 2025), somatic (Coorens, Oh et al. 2025), and culture-acquired structural (International Stem Cell, Amps et al. 2011) variants (SVs). The first is of interest to have represented in the iPSC lines, the latter two are not. The creation of clonal iPSC lines will reduce genetic heterogeneity created by somatic and reprogramming events. Therefore, we derived clonal iPSC sublines from the iPSC pools by low-density plating of dissociated single cells. Colonies were picked, expanded in StemFlex media, and archived to create a Master stock (32 vials, passage 5). During this expansion, we noted abnormal growth and morphology in iPSC sublines derived from three individuals (GeDiY151, GeDiY620, GeDiX868). These lines were also later found to differentiate poorly, prompting us to exclude them from the current GeDi panel.

To identify and exclude iPSC clones that contain rare somatic or culture-acquired SVs, we compared three single cell-derived clones from each of the nine donors to their respective donor’s pool of PBMCs. Facilitated by SNP array data, we identified at least one subline that matched the PBMC sample and was thus devoid of novel SVs (**Table S1)**. Of the total 27 single cell-derived clones generated across the nine donors, six sublines harbored novel SVs, none of which were recurrent between sublines within the same donor or from different individuals – these sublines lines were subsequently excluded. None of the selected sublines showed an obvious copy number gain at 20q, the region containing BCL2L1 that, when expanded, confers a growth advantage (Avery *et al. Stem Cell Reports* 2013). Interestingly, in some sublines we detected SV losses at 7p14.1 and 14q11.2 — regions containing T cell receptor genes (**Table S1**), suggesting that the reprogrammed cell of origin was not an erythroid progenitor as intended, but rather a circulating T cell.

To confirm clearance of the Sendai virus vector, the nine GeDi sublines were stained at passage 7 for the viral haemagglutinin-neuraminidase (HN) protein and analyzed by flow cytometry (**Figure S1A**). All nine iPSC lines showed near-complete Sendai virus clearance (<1% HN-AF647+), in contrast to cells from an early passage iPSC pool that showed 70% HN-AF647+ cells and served as a positive control for this assay. To assess chromosome stability during extended culture, G-band karyotyping was performed for each line at passage 20. No chromosomal gains, losses, or rearrangements were observed in a sampling of 20 nuclei (data not shown), indicating that the nine GeDi sublines are able to maintain euploidy after prolonged passaging.

### WGS-based variant screening identifies a pathogenic BCOR mutation in one subline

To identify mutations of clinical significance and recurrent mutations in genes known to confer a growth advantage in iPSCs, Illumina short-read sequencing was performed on the nine GeDi lines, and variant calls (DRAGEN) were annotated using Illumina Emedgene software. In addition to variants in the ClinVar database, we looked for disruptive variants in a defined list of cancer genes, including *TP53* and *BCOR*, which are commonly mutated in human PSCs (**Table S2**). Private to each of the GeDi lines, a small number of likely pathogenic mutations were identified (**Table S2**). This is not an unexpected outcome when deeply profiling human iPSCs by WGS (Hildebrandt, Reuter et al. 2019). The origin of these mutations, whether present in the original donor or culture-acquired, was not addressed here.

All but one of the GeDi lines lacked mutations in cancer genes (**Table S2**). In GeDiY518.1J, Emedgene variant annotation identified a splice donor variant in *BCOR* (GT to GC; chrX:40,054,254; **Figure S2A**) and was predicted to cause mis-splicing of the transcript, most likely leading to a truncation of the protein. Indeed, RNA-seq of undifferentiated iPSCs indicated reads extending into the second-to-last intron of this gene in GeDiY518.1J, not seen in any of the other GeDi lines (data not shown) but present in all three sub-clones derived from the iPSC pool of this donor. Loss-of-function mutations in *BCOR*, a transcriptional repressor encoded on the X chromosome, are common in human iPSCs, particularly in cell lines reprogrammed from blood (Rouhani, Zou et al. 2022) and are speculated to drive proliferation in culture.

Thus, we assessed proliferative capacity across all nine GeDi lines and KOLF2.1J by IncuCyte live-cell imaging over 84 hours (**Figure 1B; Figure S1B**). Doubling times ranged from 23.6 h (GeDiY518.1J) to 32.3 h (KOLF2.1J) and were reproducible across three independent experiments, with statistically significant pairwise differences confirmed by one-way ANOVA with Tukey post hoc correction (**Figure S1C**). This line-intrinsic variation likely reflects distinct genetic backgrounds across donors, with culture conditions (vitronectin coating, StemFlex medium) held constant across all measurements, and suggests BCOR mutant iPSCs have a faster growth rate.

To understand the functional consequence of this BCOR variant in more detail, we generated an isogenic corrected line (GeDiY518.1.1J) by CRISPR prime editing (**Figure S2B**). Compared to the corrected line, the BCOR mutant showed markedly altered colony morphology (**Figure S2C**) and a distinct transcriptomic profile, clustering separately in PCA from all other iPSC lines, including its isogenic counterpart (**Figure S2D-E**). While this BCOR mutant line differentiated to all tested lineages (**Figure S2F-J**), in all cases it remained a transcriptional outlier relative to the BCOR wt lines (**Figure S2**). Across lineages, the magnitude of transcriptional divergence was inversely correlated with BCOR expression levels in the mutant relative to wildtype, with the greatest number of DEGs observed in lineages showing the largest reduction in BCOR expression, such as the cortical brain organoid and extra-embryonic mesenchyme (R = −0.82, p = 0.046; **Figure S2K**).

GeDiY518.1J’s distinct transcriptomic profile and increased proliferation rate together support the interpretation that loss of BCOR function confers a proliferative growth advantage in iPSCs, consistent with the strong positive selection pressure for BCOR mutations documented across large iPSC repositories, where such mutations are found in 26.9% of blood-derived iPSC lines and are associated with functional transcriptomic consequences (Rouhani, Zou et al. 2022). Given the strong effect of the BCOR variant on transcriptomic state and proliferative behavior, GeDiY518.1J was excluded from all subsequent analyses, and only eight GeDi lines were carried forward for further characterization.

### The eight-line GeDi panel retains intact p53 function and is compatible with standardized CRISPR editing workflows

We next assessed whether the remaining eight-line panel retained functional p53 pathway integrity, as p53 is another pathway recurrently disrupted during iPSC reprogramming and culture (Merkle, Ghosh et al. 2017). We used a stable mCherry reporter driven by 13 tandem p53 response elements (PG13-mCherry), delivered by piggyBac transposition and adapted from the assay described for the KOLF2.1J reference line (Pantazis, Yang et al. 2022). Under basal conditions, all lines showed low mCherry expression (3.0–30.3%), confirming minimal constitutive p53 activity. Following treatment with doxorubicin to induce DNA double-strand breaks, the KOLF2.1J control cell line showed robust mCherry induction (63%), consistent with a functional p53 response. In contrast, a KOLF2-derived TP53 homozygous KO iPSC line created for MorPhiC (Adli, Przybyla et al. 2025) showed minimal mCherry expression (2%) and no response to doxorubicin treatment, confirming assay specificity All eight GeDi lines, together with KOLF2.1J, showed a marked increase in mCherry+ cells upon doxorubicin treatment (50– 70%), demonstrating intact p53-mediated DNA damage signaling across the full panel. (**Figure 1C**).

In subsequent analyses, the selected eight GeDi lines were compared with two isogenic reference lines: KOLF2.1J (Pantazis, Yang et al. 2022) and KOLF2.2J, a derivative of KOLF2.1J where a single edit was made to correct a previously identified heterozygous point mutation affecting *COL3A1* in KOLF2.1J (Hildebrandt, Reuter et al. 2019) (**Figure S1D**). Mutations in *COL3A1* are causative of Ehlers–Danlos Syndrome (Malfait, Francomano et al. 2017), and this mutation creates an alternate splice acceptor that generates a transcript with an alternate reading frame that terminates within exon 48 of a 51-exon gene, leading to nonsense-mediated decay and significantly reduced expression from the mutated allele. The corrected COL3A1 is expressed at approximately twofold the level of the parental line in KOLF2.2J-derived extra-embryonic mesenchyme (**Figure S1E**). KOLF2.2J is a reference line used across the MorPhiC and SSPsyGene Consortia. This corrected line, along with KOLF2.1J, provided the isogenic reference background used for downstream comparisons throughout this study.

To further demonstrate compatibility with standardized genome editing workflows (Skarnes, Pellegrino et al. 2019), we introduced a targeted knock-out of GCM1 — a transcriptional regulator of trophoblast differentiation (Jeyarajah, Jaju Bhattad et al. 2022) — across the eight-line GeDi panel and KOLF2.1J. Briefly, co-delivery of Cas9 ribonucleoprotein and an oligonucleotide repair template encoding a premature termination codon and two degenerate bases (termed PTC+2, a modification of PTC+1; Adli, Przybyla et al. 2025; Pellegrino, McDonough et al. 2025), was used to introduce a stop codon and frameshift in exon 2 (ENSE00000929857) of GCM1 (**Figure S1F**). The addition of degenerate bases provides a convenient barcoding strategy to identify non-sibling clones with correct biallelic modification by Sanger sequencing. Sixteen clones per line were picked and genotyped, and homozygous knock-out clones were recovered for every line, confirming that efficient biallelic editing is readily achievable in all GeDi lines (**Figure S1G**).

### Genetic analysis of GeDi lines displays population diversity in cohort

To evaluate whether our donor selection strategy, which relied on self-reported ancestry, yielded a panel spanning a broad range of genetic diversity, we projected SNP-array genotypes from the GeDi iPSC lines onto principal components derived from the 1000 Genomes Project (1KG) reference panel (Genomes Project, Auton et al. 2015). We also assessed where the lines positioned relative to regions of genetic space sparsely represented by 1KG samples. As a secondary, technical check on sample identity, we confirmed that each line’s genetic similarity to 1KG reference populations was concordant with the corresponding donor’s self-reported ancestry. Projection onto PC1 and PC3 (49.35% and 7.87% of the variance captured by the retained components) (**Figure 1D**), resolved all five 1KG reference groups into distinct regions and separated the GeDi lines more clearly than PC1 and PC2 (26.85%) (**Figure S3A**), on which the 1KG-SAS-like and 1KG-AMR-like samples overlap along the cline between the 1KG-EUR-like and 1KG-EAS-like clusters. Notably, GeDiY030 falls within the 1KG-AMR/SAS-like cluster on PC1 and PC2 but is displaced from all reference groups on PC3, consistent with the absence of Middle Eastern populations from the 1KG panel.

GeDi line similarity to 1KG reference populations matched donor self-report in every case. The two lines from donors self-reporting African (AFR) ancestry (GeDiY298, GeDiX697) were 1KG-AFR-like; the line from the donor self-reporting East Asian (EAS) ancestry (GeDiY634) was 1KG-EAS-like; the reference line KOLF2.1J, whose donor reported European (EUR) ancestry, was 1KG-EUR-like; and the line from the donor self-reporting South Asian (SAS) ancestry (GeDiX002) was 1KG-SAS-like. The three lines from donors self-reporting Hispanic/Latino or admixed American (AMR) ancestry (GeDiX927, GeDiY016, GeDiX395) fell on or near the 1KG-AMR reference samples, consistent with similarity to multiple reference groups. The line from the donor self-reporting Middle Eastern ancestry (GeDiY030) — a region not represented by any 1KG reference group — did not overlap with any reference group. This line showed close similarity in PCA space to one of the other 23 original donors that self-reported country of origin as Iran (data not shown) suggesting GeDiY030 has close similarity to Persian populations.

Human migration has occurred across super-populations in recent centuries and has created significant mixed genetic ancestries within individuals (Hellenthal, Busby et al. 2014). This is most prominent in Latin American/Hispanic populations (Medina-Munoz, Ortega-Del Vecchyo et al. 2023), from which three of eight GeDi lines are derived. Such admixture may contribute to phenotypic variation in iPSC models of human biology and disease thus we next sought to characterize the extent of this. Utilizing supervised ADMIXTURE, we established a quantitative, per-line view of these relationships derived from the SNP array data (**Figure 1E & Table S3**). Lines from donors reporting a single ancestry (GeDiY298, GeDiY634, GeDiX927, and KOLF2.1J) were assigned almost entirely to the expected reference group. While GeDiX697 was predominantly 1KG-AFR-like, consistent with self-reporting, a minor fraction of its genome (4%) shared similarity to 1KG-EUR. The South Asian donor-derived line (GeDiX002) largely shared similarity to 1KG-SAS population (87%), though also shared 10% similarity to 1KG-EUR (Italy). GeDiY030 was modeled as a mixture of 1KG-EUR- and 1KG-SAS-like similarity, consistent with its intermediate position in the PCA and with the absence of a Middle Eastern reference group in the 1KG panel. Lines from donors reporting multiple ancestries or Hispanic/Latino ancestry showed correspondingly mixed similarity profiles. Within the 1KG-AMR reference samples, GeDiX395 had the highest similarity (39%) to the Puerto Rico sub-group We next characterized the genetic diversity of each line from its whole-genome DRAGEN call set (**Figure 1F**). Across the eight GeDi donor lines, we identified 14,027,638 total unique variants (14,610,555 when KOLF2.1J is additionally included as a reference line). Total per-line variant counts ranged from approximately 4.97 to 6.13 million. The two lines from donors of self-reported African ancestry (1KG-AFR-like; GeDiY298, GeDiX697) carried the largest number of variants (6.11M and 6.13M), the highest heterozygosity rates (0.412 and 0.415), and by a wide margin the most private (cohort-singleton) variants (1.54M and 1.45M). The remaining seven lines were comparatively similar to one another, with 4.97–5.31M total variants, heterozygosity rates of 0.293–0.357, and 0.55–0.71M private variants. The one line (GeDiX395) of these seven that did show slightly higher numbers had the greatest contribution of African ancestry (11%) based on our admixture analysis.

GeDiX927 was the least diverse line in the panel, with the smallest numbers across all three of these metrics: it carried the fewest total variants (4.97M), the lowest heterozygosity rate (0.293), and the fewest private variants (588K). It was closely followed by GeDiY634 (5.05M variants; heterozygosity rate 0.303). Notably, GeDiX927 was also the only line whose genetic similarity in the ADMIXTURE analysis loaded essentially onto a single reference population (1KG-PEL, Peruvian in Lima, Peru). The donor it is derived from self-identified American Indian with Peru as their country of origin. Its low diversity and elevated homozygosity are consistent with the founder effects and reduced genetic diversity characteristic of Indigenous-American-ancestry reference populations such as 1KG-PEL (Ramachandran, Deshpande et al. 2005, Pemberton, Absher et al. 2012, Harris, Song et al. 2018). Altogether, our genetic analyses were concordant with donor self-reported ancestry.

### Individual-specific identity markers

To support sample-identity verification across assays and data types, we generated per-line lists of private SNVs called against GRCh38/hg38 for the eight GeDi lines and KOLF2.1J. For each line we provide two sets: a genomic set, comprising all variants private to that individual within the cohort, and a transcriptomic set, restricted to private variants in transcribed exonic regions (CDS, UTR or exon) and thus potentially observable in RNA-based data. Across the panel we identified 53,312 genomic markers (3,280–14,457 per line) and 5,886 transcriptomic markers (333–1,566 per line). The number of markers mirrored the per-line diversity results; because each variant is carried by only a single individual in the cohort, these markers function as a genetic fingerprint that can be used to confirm line identity and to flag cross-contamination or sample swaps—at the whole-genome level using the genomic set, or in transcriptomic/RNA-based datasets using the transcriptomic set. The complete lists, annotated with GRCh38 position, genotype, overlapping gene and feature class, are provided as two combined tables spanning all lines (one genomic, one transcriptomic; **Dataset S1 and S2**, respectively).

### Multi-lineage differentiation potential of GeDi lines

To assess whether the differentiation protocols produced consistent, lineage-specific transcriptomes across the eight GeDi iPSC lines, we differentiated these lines and the KOLF2.2J reference down multiple lineages. Monolayer cultures of primitive syncytium (PrS) and extra-embryonic mesenchyme (ExM) through 6 days of differentiation represented extra-embryonic lineages (Liu, Tan et al. 2025). Monolayer cultures of endoderm (Endo) and mesoderm (Meso) leveraged commercially available protocols, and were differentiated through 5 days. For representation of the ectoderm lineage, we generated very early stage (Day 9) cortical brain organoids (CBO). Bulk RNA-seq was generated in biological triplicate at the end of each differentiation, and for the undifferentiated cells (Und). We computed Euclidean distances between all samples on variance-stabilized expression and clustered them hierarchically (**Figure 2A**). Samples partitioned cleanly by lineage rather than by individual or batch: undifferentiated samples formed a tight block, and each differentiated lineage resolved into its own cluster. This lineage-dominant structure confirms both that the undifferentiated lines maintain a consistent pluripotent identity and that the differentiation schemes are robust and reproducible across all donors — replicates within each lineage were highly concordant regardless of genetic background. The numbers of significantly up-regulated genes per lineage were 1,600, 2,196, 1,674, 3,223, and 3,595 for CBO, Endo, Meso, ExM, and PrS, respectively. The panel of eight GeDi lines also expressed key master regulators consistent with each differentiated lineage (see Methods), making it a resource for both embryonic and extra-embryonic cell fates.

**Figure 2.**
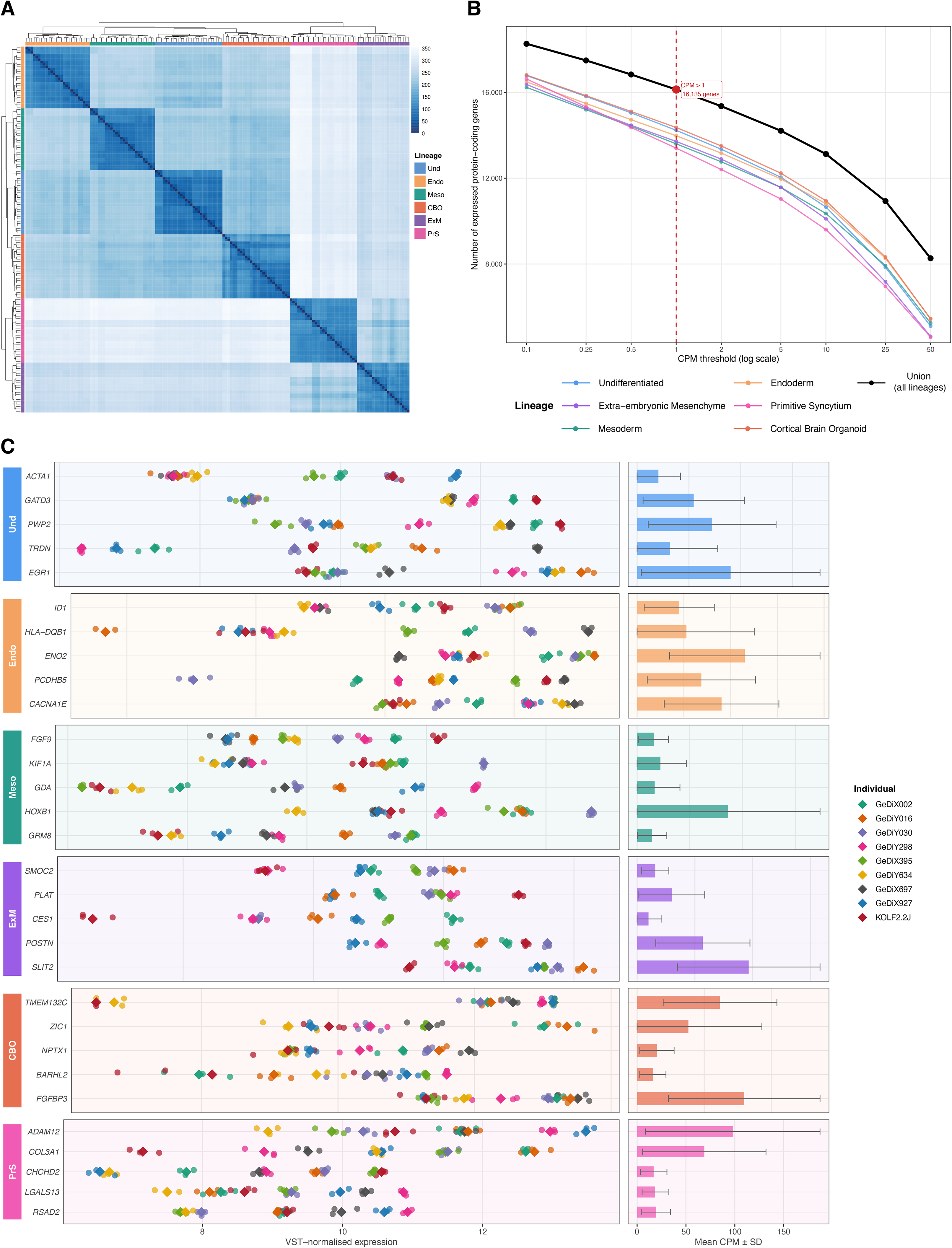
Transcriptomic characterization of the GeDi iPSC panel across undifferentiated and differentiated states. Nine iPSC lines (eight GeDi lines and the KOLF2.2J reference) were profiled by total RNA-seq in the undifferentiated state (Und) and across five lineages: definitive endoderm (Endo), mesoderm (Meso), dorsal-forebrain cortical organoid (CBO), extra-embryonic mesenchyme (ExM) and primitive syncytium (PrS). **(A)** Heatmap of sample-to-sample Euclidean distances, with samples ordered by hierarchical clustering and identical row and column ordering. Color indicates distance, from dark blue (most similar) to white (most distant). The color bars at the top and left annotate the lineage of each sample. **(B)** Sensitivity of expressed protein-coding gene detection to the CPM threshold. For a range of CPM thresholds (0.1–50, log scale), the number of expressed protein-coding genes is shown for each lineage and for their union across all lineages (black). The vertical dashed line marks a standard threshold (CPM > 1), at which the union comprises 16,135 protein-coding genes (80.2% of the 20,121 protein-coding genes annotated in GENCODE v49). **(C)** Five representative highly variable genes across donors, shown per lineage, grouped in lineage-colored blocks (labelled at left, gene symbols at right of each block). Left: VST-normalized expression, where each small point is a replicate and each large diamond a per-individual mean. Right: mean CPM (± SD, error bars) for each gene across the lineage.

Each differentiated lineage showed the expected canonical marker signature, including definitive endoderm and primitive-streak regulators in Endo, paraxial-mesoderm and HOX genes in Meso, neuroectodermal/forebrain markers in CBO, mesenchymal/ECM genes in ExM, and a canonical PrS signature (including CGA, the most abundantly expressed protein-coding gene in PrS across all lines) (representative markers listed in **Table S4**; full differential expression results in **Dataset S3**). The reciprocal contrast (undifferentiated vs. all differentiated samples) yielded 963 up-regulated genes constituting a clear pluripotency signature (*POU5F1*, *NANOG*, *PRDM14*, and others), confirming that undifferentiated lines retain a robust pluripotent identity that is appropriately extinguished upon differentiation. Together, these results demonstrate that the GeDi panel supports faithful differentiation across both embryonic and extra-embryonic fates.

To gauge how comprehensively the differentiation panel samples the human transcriptome, we quantified the number of expressed protein-coding genes (GENCODE v49) per lineage and across their union, as a function of the CPM threshold used to call a gene expressed (**Figure 2B**). At the chosen threshold of CPM > 1, the union across all lineages comprised 16,135 protein-coding genes — 80.2% of the 20,121 protein-coding genes annotated in GENCODE v49. Thus, across embryonic and extra-embryonic fates the panel expresses a large and stable fraction of the protein-coding genome despite representing only very early stages of differentiation (5 to 9 days).

### Highly variable genes reveal functional diversity across donors

Because the central aim of this panel is to capture genetic and functional variation between lines, we identified the genes most variable between individuals within each lineage, restricting to autosomal genes and combining a likelihood-ratio test with a dispersion-based (MAD) ranking. Every lineage contained genes that were both well expressed and strongly variable across donors. The five representative variable genes shown per lineage (**Figure 2C**) are expressed at appreciable levels (mean CPM) yet differ markedly in magnitude between individuals, with per-individual means (diamonds) spread across the expression range while replicates of the same donor remain tightly grouped — indicating that the variability reflects stable, donor-intrinsic differences rather than technical noise. Representative examples include *ACTA1*, *PWP2*, *EGR1* (Und); *ID1*, *HLA-DQB1*, *ENO2*, *CACNA1E* (Endo); *KIF1A*, *GDA*, *HOXB1*, *GRM8* (Meso); *SMOC2*, *PLAT*, *POSTN*, *SLIT2* (ExM); *TMEM132C*, *ZIC1*, *NPTX1*, *BARHL2*, *FGFBP3* (CBO); and *ADAM12*, *LGALS13*, *RSAD2* (PrS). This donor-dependent variability, observed even within transcriptionally well-defined lineages, underscores the value of the panel for studying how diverse genetic backgrounds shape lineage-specific gene expression.

### Haplotype-resolved genome assemblies and structural characterization of the GeDi panel

Short-read whole-genome sequencing, while powerful for SNV and small indel detection, is fundamentally limited in its ability to resolve variants within repetitive regions and complex genomic loci. Specifically, the resolution of genetic variation within segmentally duplicated regions, tandem repeats, and even mobile elements is inhibited by the inability to uniquely map short reads in these loci (Logsdon, Ebert et al. 2025). To generate a comprehensive catalog of genetic variation across the GeDi lines, we produced haplotype-resolved genome assemblies. For each line, we generated approximately 40× PacBio HiFi and 20× ultra-long ONT coverage (reads ≥100 kb), along with Dovetail Hi-C for phasing. Using these data, we produced highly contiguous haplotype-resolved assemblies for the eight GeDi lines and KOLF2.1J with hifiasm. The assemblies achieved a median area under the Nx curve (auN) of 128.6 Mb (**Table S5, Figure S4E**), with a median of 78 contigs per haplotype assembly (range 59–109) and 87% of contigs exceeding 50 kb with an average QV value of 64 estimated by k-mer analysis (Figure S4F).

Assembly metrics were consistent with current approaches (Table S5) (Logsdon, Ebert et al. 2025) (Huang and Li 2023, Tegenfeldt, Kuznetsov et al. 2025). Assemblies containing the Y chromosome showed slightly lower completeness (median 97.2%), likely reflecting the reduced representation of Y-linked orthologues in BUSCO mammalian gene sets rather than reduced assembly quality. Assessing assembly accuracy with Flagger, we estimated that 99.2% of the phased assembly sequence was correctly assembled (**Figure S4G**). Alignment to GRCh38 showed a median genome coverage of 95.6% (Figure 3B and **Table S5**), and most chromosomes were represented by a single contig in at least one haplotype, chromosome Y being the primary exception due to the lack of presence in female lines (**Figure 3A**).

**Figure 3.**
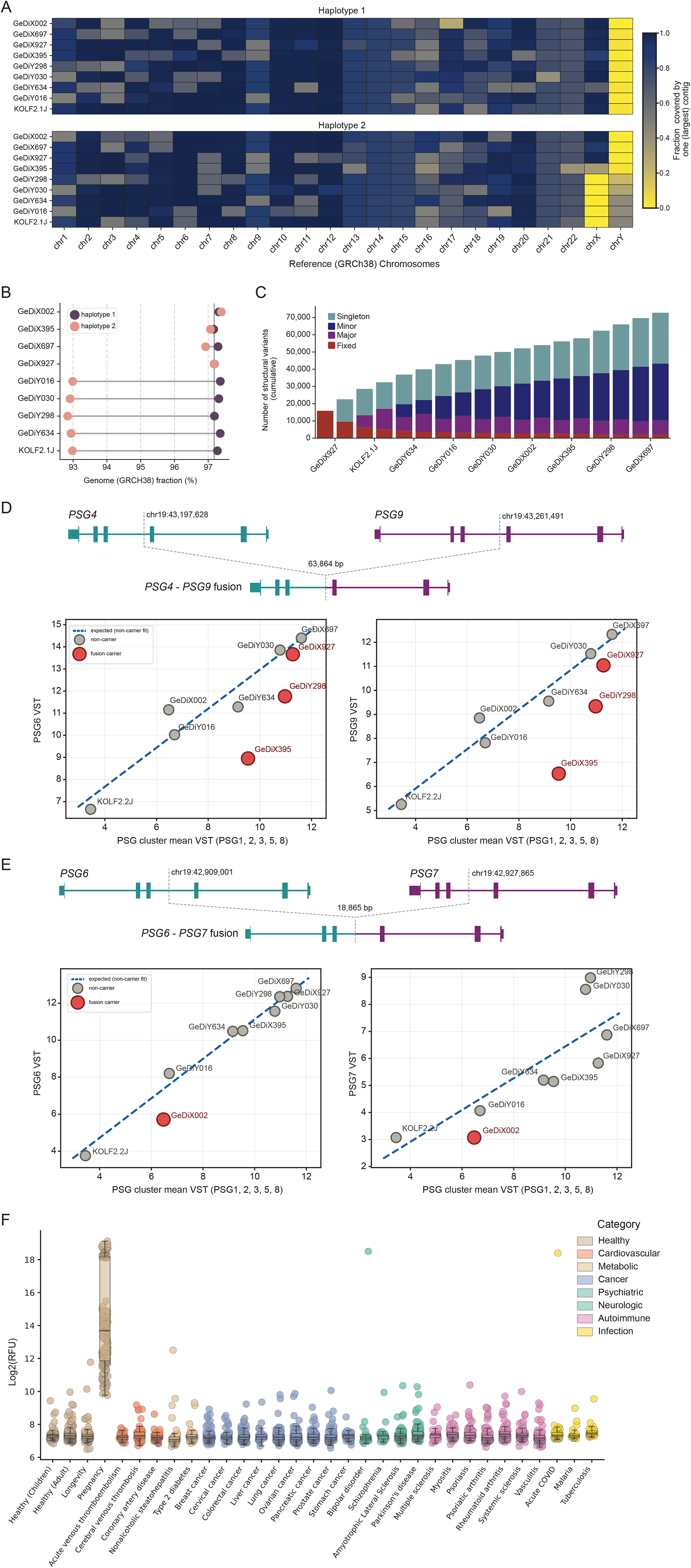
Genome assembly quality and structural variation in iPSC lines. Haplotype-resolved (h1, h2) T2T assemblies were generated for nine iPSC lines and used to characterize structural variation at the PSG cluster (chr19q13.31). **(A)** Assembly contiguity, measured as the fraction of each GRCh38 reference chromosomes spanned by the largest contigs, for each haplotype in all cell lines. Reduced chrX and chrY coverage reflects sex chromosome complement of the corresponding lines. Haplotypes containing chrY were designated as haplotype (hap2). **(B)** Genome fraction (%) of GRCh38 per haplotype across all cell lines. Haplotypes containing chrY were designated as haplotype (hap2). **(C)** Cumulative non-redundant SVs identified across all haplotypes relative to GRCh38. **(D)** Structural variant at the PSG4/PSG9 locus and its expression consequences. Top: gene models for PSG4 and PSG9 (boxes, exons) with deletion breakpoints (chr19:43,197,628 and chr19:43,261,491, hg38); the intervening ∼63.9 kb is deleted, producing the PSG4–PSG9 fusion (bottom track). Bottom: expression of PSG4 and PSG9 under primitive-syncytium (PrS) differentiation, plotted against the PSG-cluster reference (per-line mean variance-stabilized expression (VST) of the structurally intact cluster genes PSG1, 2, 3, 5 and 8) across the nine lines. The dashed line is the expectation fit on non-carrier lines; grey points are non-carriers and red points are fusion carriers (GeDiX395, GeDiX927, GeDiY298). Carriers fall below expectation for both genes. **(E)** As in (D) for the PSG6/PSG7 locus: breakpoints chr19:42,909,001 and chr19:42,927,865 delete ∼18.9 kb to produce the PSG6–PSG7 fusion, and the carrier (GeDiX002) is under-expressed for both genes. **(F)** Circulating PSG7 levels vary widely in pregnancy. Box-and-jitter plot of PSG7 relative abundance (log2 RFU, SomaScan) across 27 healthy and disease cohorts from the Human Protein Atlas pan-disease blood proteome resource (n = 2,356 samples total). Each point represents one individual; boxes show the median and interquartile range, colored by physiological/disease category. While PSG7 levels are tightly distributed and largely overlapping across non-pregnant states — regardless of disease category — pregnancy samples show markedly elevated levels spanning nearly 10 log2-units.

The resultant assemblies enable allelic analysis of SVs and regions of genomic complexity that have historically been refractory to characterization in non-lymphoblastoid origin cell lines (Liao, Asri et al. 2023, Lucas, Hebbar et al. 2026). Notably, more than 98% of the 22q11.21 locus was spanned by a single contig across assemblies (**Figure S4B**), enabling precise characterization of structural variation within this complex medically relevant locus known to harbor rearrangements resulting in DiGeorge syndrome. Moreover, our contiguous assemblies spanned existing gaps in the hg38 20q11.21 locus, resolving both haplotypes in each cell line and enabling variant calling within a complex locus that is frequently amplified in human iPSC lines (International Stem Cell, Amps et al. 2011), and human cancers (Krivec, Ghosh et al. 2024).

### Variant calling

Across the nine cell lines, we identified 13,360,800 non-redundant variants, comprising 10,590,802 single-nucleotide variants (SNVs), 2,697,229 small insertions and deletions (indels; 1,344,056 insertion 1,353,173 deletions) and 72,769 structural variants (SVs; 45,314 insertions, 27,394 SV deletions, and 61 inversions) (Figure 3C, S4H, S5I and Table S9). Collectively these variants spanned 10.6 Mb of the GRCh38 reference genome for SNVs, 11.2 Mb for indels, and 67.8 Mb for SVs. SVs insertion accounted for the largest cumulative span (35.8 Mb), highlighting the contribution of sequences absent from GRCh38 reference genome. The median variant counts per cell line were 3,942,851 single-nucleotide variants (SNVs), 968,868 small insertions and deletions (indels; 483,346 deletions and 485,522 insertions), and 23,351 structural variants (SVs; 8,970 deletions, 14,359 insertions, and 22 inversions). Median SVs sizes were 130bp for deletions, 176bp for insertions and 8,717 bp for inversion. We further categorized 2,138 variants (with a median of 238 variants per cell line (Table S10)) as high-priority variants (Methods) by variant effect predictor.

### Structural variation in the PSG cluster alters expression of the rearranged genes

The Pregnancy Specific Glycoprotein (*PSG*) genes, expressed in the PrS RNA-seq data, were among the most variably expressed across the GeDi lines (**Dataset S4**). The PSG locus is rapidly evolving with 11 in tandem copies in the human genome (Moore, Williams et al. 2022) and with a high frequency of SV and gene conversion-associated polymorphisms within the human population (Chang, Semyonov et al. 2013, Dumont and Eichler 2013). PSGs are specifically expressed in the trophoblast syncytium and are among the most abundant fetal-derived proteins in maternal circulation where they appear to stimulate an anti-inflammatory phenotype on monocytes (Snyder, Wessner et al. 2001). Levels of PSGs in maternal circulation have also been linked to preeclampsia (Kandel, MacDonald et al. 2022). With the expression variation across GeDi lines and the known association between this locus and disease, we investigated our assemblies for structural variation in PSGs.

It is unlikely the variation in expression identified for the PSG genes within the PrS is derived from line differences in states of differentiation as mRNA for other PrS-derived secreted proteins detectable in maternal serum (CGA, GDF15, INSL4, and PGF) do not vary to a similar extent (**Dataset S3**). There were no obvious DNA methylation differences within this locus across lines (data not shown). Thus, the PSG expression differences likely reflect underlying genetic variability. We identified numerous deletion rearrangements, and the genes disrupted by each deletion were specifically under-expressed in carriers (**Figure 3D-E**). Heterozygous carriers of the *PSG9–PSG4* fusion fell below expectation by 2.04 VST for *PSG4* and 2.36 VST for *PSG9*, and the *PSG7–PSG6* fusion carrier (GeDiX002) fell below by 1.65 VST for *PSG6* and 1.30 VST for *PSG7* — the direction predicted by the allelic loss of one promoter and canonical gene body. The effect was confined to the rearranged genes; the remaining cluster members reflected gene expression that fell in the within-cluster reference. In addition to these SVs, we also identified a CGA→TGA transition at PSG7 codon 98 (p.R98*) in 16 of 18 haplotypes, biallelic in seven of nine lines and absent only from GeDiY030-h1 and GeDiY298-h2. This is the same nonsense variant recently described as a common polymorphic pseudogenization of PSG7 (Lopes-Marques, Peixoto et al. 2024). Thus, within our 8 GeDi lines and KOLF2.1J we observe substantial variation within this locus, resulting in variable expression of PSGs across lines, which may reflect the measured differences of PSGs in maternal blood identified in the human population (**Figure 3F**).

### Two GeDi lines carry signatures of T-lymphocyte origin

Although PBMCs were enriched for erythroid progenitor cells prior to reprogramming, two lines — GeDiY298 and GeDiY634 — carried signatures indicative of a T-lymphocyte cell of origin. We first noted T-cell receptor rearrangements from SV calls from the SNP array data for these lines (discussed above), and both lines expressed markedly lower levels of the T-cell receptor γ and δ constant-region genes *TRGC2* and *TRDC* than the rest of the panel in the analysis of RNA-seq of the derived mesoderm lineage cells (**Dataset S4**). Inspection of the genome assembly-based phased assembly variant (PAV) calls revealed heterozygous deletions within the T-cell receptor β (*TRB*) locus at chr7q34 — 242 kb in GeDiY298 and 325 kb in GeDiY634 — that removed the trypsinogen genes *PRSS1* and *PRSS2* interspersed among the TRBV segments, the expected genomic footprint of a somatic TRB V(D)J rearrangement. We found no cell-of-origin effect on methylation, imprinting, or allele-specific expression (ASE) for these two T-lymphocyte origin lines.

### Genomic methylation status across GeDi lines

We leveraged the DNA methylation data captured by long-read sequencing (ONT) to analyze global methylation states across the GeDi lines. We first measured methylation across major genomic compartments. All lines showed the expected partitioning. Methylation increased progressively across the CpG-island neighborhood: CpG islands were largely unmethylated (mean 17.2%, range 15.9–18.6%) and gained methylation with distance from the island, rising through the flanking CpG shores (61.2%) to the more distal CpG shelves (80.7%). Promoters were similarly hypomethylated (27.0%, 26.5–27.6%), whereas gene bodies (75.2%) and repetitive elements (LINE 84.3%, SINE 87.3%, LTR 82.9%) were highly methylated, near the genome-wide average of 77.0% (75.8–78.8%). These distributions match the expectation for a normal pluripotent methylome (Lister, Pelizzola et al. 2009, Jones 2012), and the across-line variation was small for every feature (genome-wide range < 3 percentage points; **Figure S5A**), indicating that global methylation is both correctly patterned and reproducible across the panel.

Global genomic methylation patterns across the GeDi lines compared favorably to other pluripotent lines and were distinct from the differentiated lineages from which they were derived. Principal component analysis of 10 kb-windowed genome-wide methylation (**Figure 4C**) positioned all eight iPSC lines as a single tight cluster (PC1 43.8% of variance, PC3 8.8%) together with embryonic stem cell (ESC; H1, H9, HUES64) and iPSC (KOLF2.1J, DF 6.9, DF 19.11) references and clearly separated from the blood references (CD14+ monocytes, CD3+ T cells, NK cells, HSC CD34+) and from somatic tissue and fibroblast methylomes (liver, lung, spleen, IMR90). Shown is PC1 versus PC3 because PC2 was dominated by assay platform (**Figure S3B-C**)—long-read ONT for our lines versus Illumina whole-genome bisulfite sequencing for the public references—and therefore separated samples by technology rather than by biology. The convergence of the reprogrammed lines onto the pluripotent reference cloud, and their displacement from blood, is consistent with successful resetting of the somatic methylome toward a pluripotent state.

**Figure 4.**
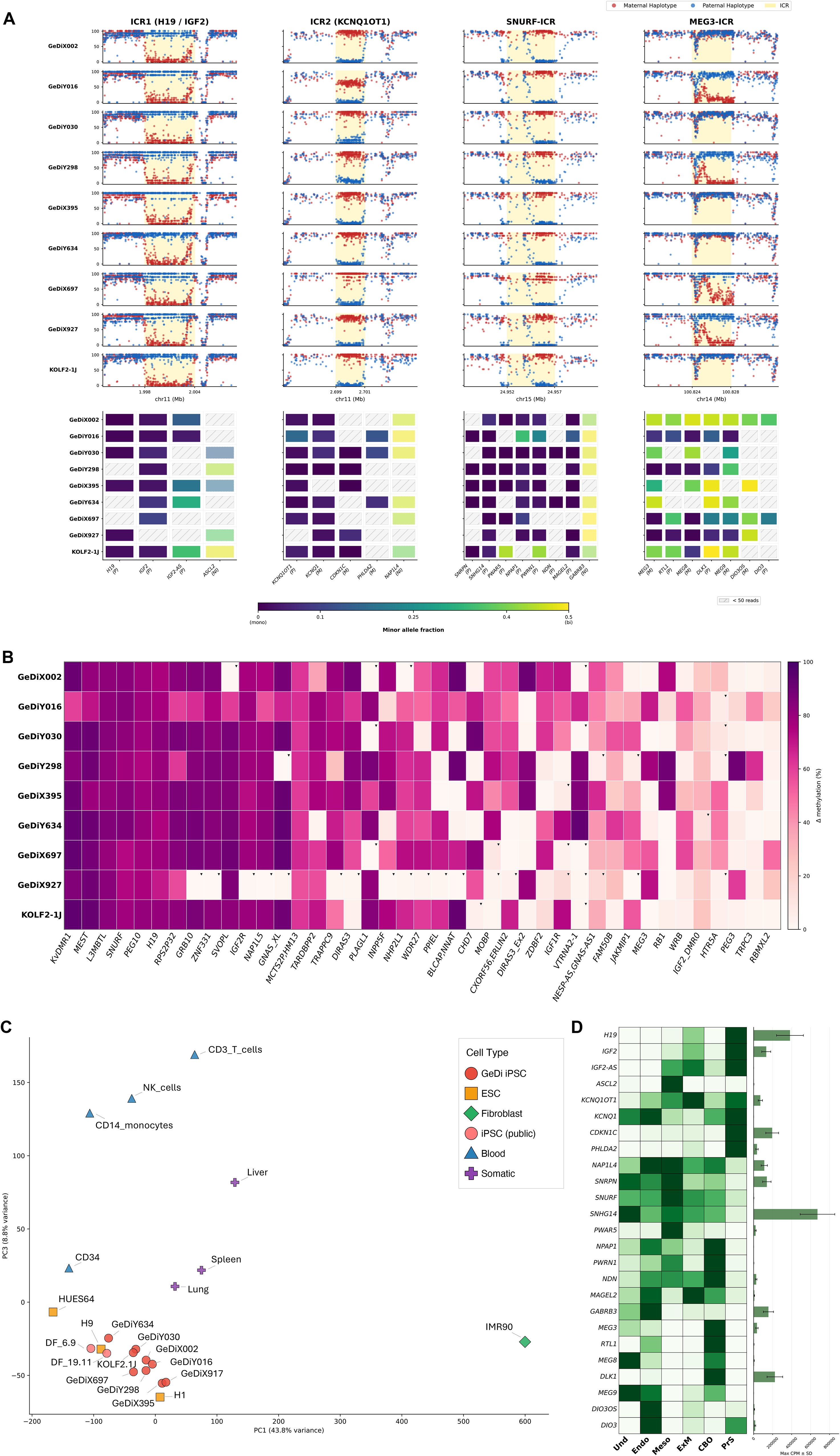
Preservation of genomic imprinting and lineage-resolved expression of imprinted loci across reprogrammed iPSC lines. Panels show data for the nine iPSC lines (eight GeDi lines and the KOLF2-1J reference). **(A)** Haplotype-resolved DNA methylation and allele-specific expression at four imprinting control regions (ICRs): ICR1 (*H19*/*IGF2*, chr11), ICR2 (KvDMR1/*KCNQ1OT1*, chr11), the SNURF/SNRPN-ICR (chr15), and the MEG3-ICR (chr14). *Top*, per-CpG methylation across each ICR ± 5 kb for every line (rows); points are colored by inferred parental haplotype (maternal, red; paternal, blue) and the ICR interval is shaded gold. The y-axis is methylation (0–100%) and the x-axis is genomic position (Mb). *Bottom*, allele-specific expression of each ICR’s target genes together with selected neighboring non-imprinted (NI) control genes; each cell is colored by the gene’s minor-allele fraction on a continuous scale from 0 (monoallelic) to 0.5 (biallelic). Gene labels indicate the parent-of-origin of expression (M, maternally expressed; P, paternally expressed; NI, non-imprinted); grey hatched cells denote genes with < 50 informative reads. **(B)** Allelic methylation difference (Δ = |hap1 − hap2|, %) at 41 germline imprinted DMRs (columns) across the nine lines (rows). Triangles mark cells classified as hypo/hypo (Δ < 20% and mean methylation < 30%), indicating candidate loss of the differential methylation imprint. **(C)** Principal component analysis of genome-wide CpG methylation (10 kb windows) for the nine iPSC lines together with public reference methylomes, shown as PC1 (43.8% of variance) versus PC3 (8.8%). Symbols are colored and shaped by cell type (GeDi iPSC, ESC, public iPSC, fibroblast, blood, somatic). **(D)** Expression of imprinted DMR target genes (rows) across six differentiation lineages (columns: Undifferentiated, Endoderm, Mesoderm, Extra-Embryonic Mesenchyme, Cortical Brain Organoid, Primitive Syncytium). Heatmap color shows the median CPM per lineage, scaled independently for each gene. The bars at right show each gene’s maximum CPM (± SD across lineages) to convey absolute expression level.

### Variable stability of methylation imprinting

As genomic imprinting controls the dosage of developmentally regulated genes, and disruption of this control for specific genes can lead to embryonic and/or perinatal lethality (Kono, Obata et al. 2004), it is important to quantify the status of these loci in reference iPSC lines. We examined haplotype-resolved methylation at 41 previously defined germline imprinted differentially methylated regions (DMRs) (Court, Tayama et al. 2014, Akbari, Garant et al. 2022, Jima, Skaar et al. 2022). The coordinates of these DMRs coincided with the differentially methylated intervals we identified directly by manual inspection of the haplotype tracks. Visual examples of such data are provided for the *NNAT* and *PEG10* DMRs, examples of unstable and stable imprints across lines, respectively (**Figure S5B-C**). A representative summary of methylation status for all DMRs associated with imprinting loci, in order of most to least stable across lines, is provided (**Figure 4B**).

The functional consequence of imprinting is monoallelic expression of genes within the neighborhood of each DMR, and with our haplotype-resolved assemblies we are able to link inferred parent-of-origin imprints to parent-of-origin monoallelic expression, at least for informative genes that contain heterozygous SNPs within their mRNA, and are expressed in our dataset. Representative examples of imprinted DMRs are shown for ICR1 (H19/IGF2), ICR2 (KvDMR1), and the SNURF/SNRPN-ICR which are preserved across all lines, together with the MEG3-ICR as a representative variable locus (**Figure 4A**). These DMRs are paired with the allele-specific expression of target genes, with selected neighboring non-imprinted genes included as controls. Allele-specific expression for genes associated with all 41 DMRs can be found in the allele-specific expression table (**Table S6**). Allele-specific expression recapitulated the methylation findings at the expression level. Of the 12,000–16,600 genes testable per line (≥ 50 informative reads), only ∼7–10% were classified as monoallelic genome-wide, and this set recovered the canonical imprinted targets.

For stable imprints, the maintenance of monoallelic expression was tight, for instance, minor-allele fractions neared zero even at very high coverage, for example, in GeDiX395, SNRPN showed a minor-allele fraction of 0.002 over 57,256 reads and SNHG14 0.003 over 130,565 reads. This is in contrast to other more variable imprinting loci across GeDi lines, for instance, at the MEG3-ICR (**Figure 4A)**. Lines that retained MEG3-ICR imprints (GeDiY016, GeDiY298, GeDiX697, GeDiX927; Δ ≈ 54–77%) showed monoallelic *MEG3* expression (minor-allele fraction 0.003–0.076), whereas lines in which the MEG3-ICR had lost its allelic difference (GeDiX002, GeDiY030, GeDiX395, GeDiY634, KOLF2.1J; Δ ≈ 0–5%) expressed *MEG3* biallelically (0.32–0.46). The concordance was essentially one-to-one across the GeDi lines, and the broader 14q32 cluster (e.g., *DLK1*) followed the same direction. This coupling between the epigenetic state of the MEG3-ICR and the allelic output of its targets illustrates the regulatory control exerted by this DMR and explains the variability in expression seen for the genes within this locus.

Genes under the control of imprinting are developmentally regulated and this developmental, lineage-restricted dynamic is captured in our lineage-specific RNA-seq data (**Figure 4D**). Notably, the ICR1 and ICR2 cluster genes (*H19*, *IGF2*, *IGF2-AS*, *KCNQ1OT1*, *KCNQ1*, *CDKN1C*, *PHLDA2*) were most strongly expressed in the extra-embryonic derivatives consistent with the known importance of these loci in extra-embryonic/placental tissues (Coan, Burton et al. 2005, Frost and Moore 2010), whereas other targets showed enrichment in neural (e.g., *DLK1*), pluripotent stem cell (e.g., *SNHG14*), endoderm (e.g., *DIO3*), and mesoderm (e.g., *PWAR5*) lineages.

### Molecular features associated with Xi erosion

X chromosome inactivation (XCI) is a fundamental epigenetic feature of female somatic cells - a form of dosage compensation leading to one active X (Xa) and one inactive X (Xi). In pluripotent stem cell cultures, the inactive X chromosome is well documented to undergo progressive erosion of inactivation over extended passage - a phenomenon referred to here as Xi erosion (Xi/e) - with different degrees of erosion occurring between lines. This erosion can alter differentiation propensities (Lessing, Anguera et al. 2013, Bansal, Ahern et al. 2021, Yokobayashi, Yabuta et al. 2021, Raposo, Caldas et al. 2025). This is not an all-or-none phenomenon; generally there remains extensive monoallelic expression from the Xi/e in eroded lines and heterogeneity across lines with respect to impacted genes (Raposo, Caldas et al. 2025). Thus, documentation of the extent of Xi/e is an important QC metric for reference iPSC lines. To begin to assess XCI status in the four selected female GeDiX lines, we quantified the proportion of nuclei containing H3K27me3 spots indicative of the inactive X chromosome (i.e., Barr body) by immunostaining (**Figure 5A; Figure S1I**)(Plath, Fang et al. 2003). As a control, GeDiY (male) lines showed near-zero H3K27me3 foci-positive nuclei (0.00–1.50%), as expected. At passage 4, GeDiX lines showed variable XCI maintenance: GeDiX697 displayed the highest proportion of XCI+ nuclei (85%), followed by GeDiX395 (79%), GeDiX002 (64%), and GeDiX927 (58%). Extending this analysis to passage 12 cells revealed a progressive decline in XCI+ nuclei in all lines, consistent with the known passage-dependent erosion of XCI in female iPSCs (**Figure S1H**). Variability in the extent of this was still apparent across lines, with GeDiX697 maintaining the highest level of Barr body+ nuclei at 62% at p12.

**Figure 5.**
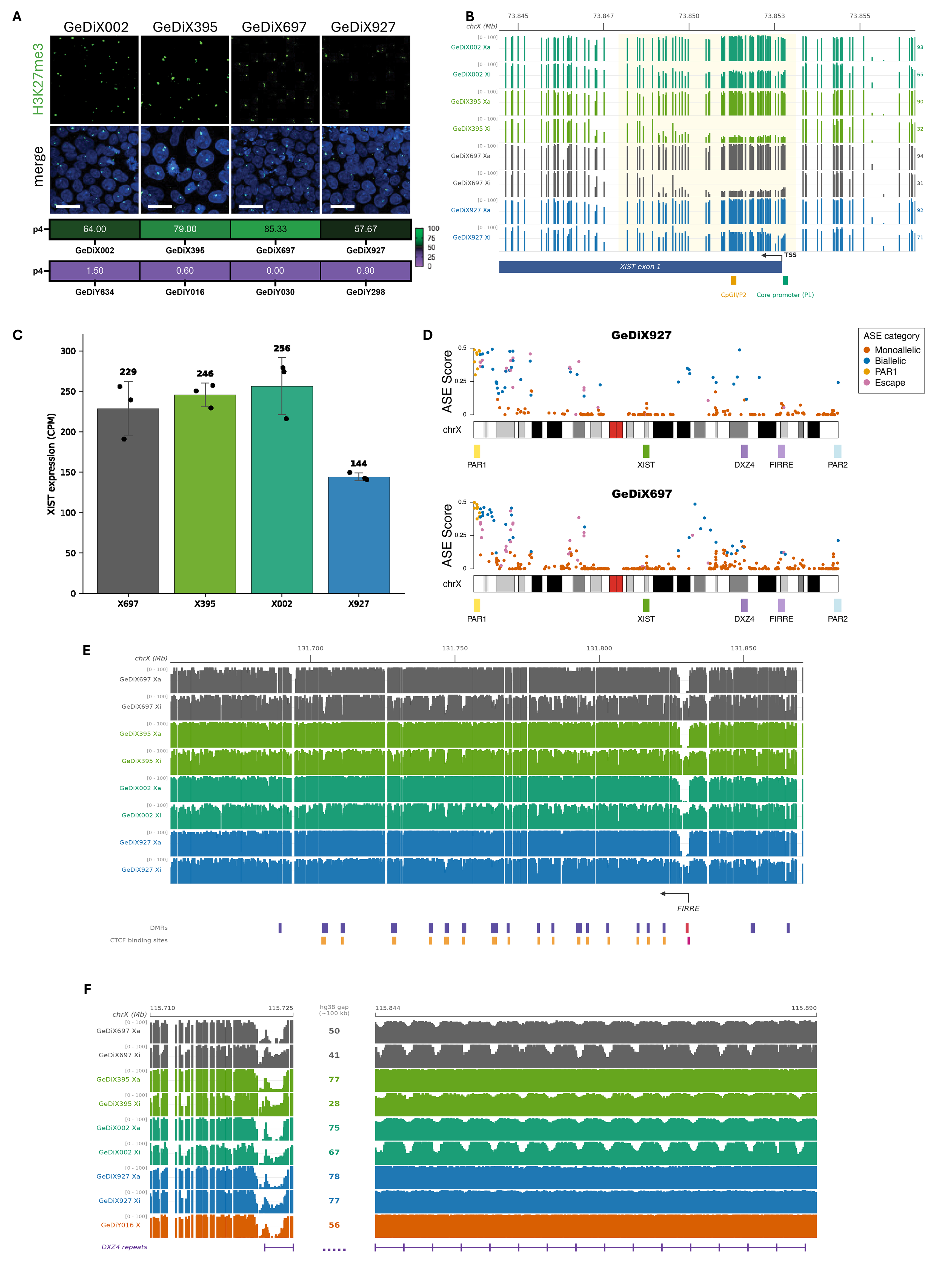
Characterization of X-chromosome inactivation erosion across GeDi iPSC lines. **(A)** Identification of condensed chromatin indicative of an inactive X chromosome by H3K27me3 immunofluorescence. Representative immunofluorescence images of the four female lines at passage 4 showing H3K27me3 (green) and merged channels (blue, DAPI). Color bar below images indicate the percentage of nuclei containing a discrete H3K27me3 focus indicative of an Xi. A similar color bar (immunofluorescence not shown) for the four male lines serves as negative controls. Scale bar = 20 μm. **(B)** Haplotype-resolved DNA methylation at the *XIST* regulatory region in the four female lines, shown as separate tracks for haplotype 1 and haplotype 2 (0–100%). Each haplotype is labeled Xa (active) or Xi (inactive) based on *XIST*-promoter methylation: the inactive X expresses *XIST* and is hypomethylated at this promoter, whereas the active X silences *XIST* and is hypermethylated. The core promoter (P1) and the exon-1 regulatory element (CpGII/P2) are marked in green and yellow, respectively, regions that are detected by array-based methylation detection methods (Bansal et al. 2021); the TSS and *XIST* exon 1 are indicated below. The full extent of the allelic differentially methylated region (DMR) recovered from the long-read assemblies is shaded in light yellow. **(C)** *XIST* expression in the four female lines (bulk RNA-seq of undifferentiated iPSCs, n = 3 each), in counts per million (CPM). Bars show the mean; points are individual replicates and error bars the standard deviation. **(D)** Allele-specific expression (ASE) across the X chromosome for (top) GeDiX927 and (bottom) GeDiX697. Each point is one informative gene (241 and 430 genes for GeDiX927 and GeDiX697, respectively), positioned at its genomic coordinate; the y-axis is the ASE score (0–0.5). Genes are colored by category: region PAR1 (amber), genes reported to escape X-chromosome inactivation (pink), biallelic (blue), monoallelic (vermillion). Locus and region markers (PAR1, *XIST*, *DXZ4*, *FIRRE*, PAR2) are shown beneath. **(E, F)** Haplotype-resolved DNA methylation across two X-linked loci, shown as separate tracks for the active (Xa) and inactive (Xi) (0–100%). Gene positions and orientation are indicated by strand-aware TSS arrows. **(E)** *FIRRE*, in the four female lines. Allelic DMRs are shown below the tracks as colored blocks: indigo, concordant with *XIST* (Xa > Xi); red, reversed (Xi > Xa). The CTCF binding sites row below the DMRs marks CTCF-bound regulatory elements (ENCODE cCREs) overlapping the DMRs: amber, CTCF-bound enhancers; magenta, the CTCF-bound promoter. **(F)** The *DXZ4* macrosatellite, in the four female lines and one male line (GeDiY016) shown for comparison. The hg38 gap (∼100 kb) is shrunken for display purposes. The purple comb marks the resolvable *DXZ4* monomers at the flanks (15 on the distal side, 1 on the proximal side), and the *total* monomer copy number per haplotype is given within the gap.

Additional molecular features of the Xi and its progressive erosion can be captured in our haplotype-resolved assembled genomes, the associated genomic methylation, and the heterozygous SNPs within the measured transcriptome. These data were generated on cells from passage 8 to 10, thus at an intermediate stage compared to the passage 4 and passage 12 Barr body quantification. The expression of the long non-coding RNA responsible for initiating XCI, X-inactive specific transcript (*XIST*), was maintained in all four lines though GeDiX927 had approximately half the expression of the other three lines suggesting this line had initiated the silencing of XIST transcription (**Figure 5C**). Increasing genomic methylation in and around the XIST promoter is an early event in XIST transcriptional silencing and Xi/e (Bansal, Ahern et al. 2021). This is reflected in the GeDiX lines where a differentially methylated region (DMR) of approximately 5 kb in length spanning the promoter and into the first exon is captured (chrX: 73,847,950-73,853,120) (**Figure 5B**). The extent of methylation here was 31%, 32%, 65% and 71% in GeDiX697, GeDiX395, GeDiX002 and GeDiX927, respectively, thus trending with the extent of Barr body maintenance. This DMR detects the greater expanse of epigenetic dynamics associated with transcriptional control of *XIST* that long-read assembly-resolved methylation provides, compared to array-based DNA methylation methods (**Figure 5B)** and corresponds to a region with the H3K4me3 histone modification known to mark promoters (as seen in ENCODE reference data).

Leveraging the heterozygosity within each line and the aggregate RNA-seq data, we next calculated allele-specific expression (ASE) along the X chromosome to determine monoallelic and biallelic expression. Total informative genes (those with heterozygosity) were 430, 347, 360, and 241 for GeDiX697, GeDiX002, GeDiX395, and GeDiX927, respectively. The percentage of biallelic genes of the total informative genes for each line was 14.89%, 23.63%, 24.38%, and 26.67%, for GeDiX697, GeDiX002, GeDiX395, GeDiX927, respectively, and included pseudoautosomal region 1 (PAR1) genes (from 9 to 13 genes per line; **Table S7**). The lines with the lowest and highest percentage of biallelic genes reflected the lines with the highest and lowest Barr body percentages, respectively, with a minor rank discrepancy for the middle two lines.

Mapping allele-specific expression across the X chromosome revealed both constitutive and erosion-associated patterns of escape from inactivation (**Figure 5D**). As expected, genes in PAR1 were biallelically expressed in all four lines (≈100% of PAR1 genes). Literature-defined escapees, genes that are not in the PARs but are known to be expressed from both Xa and Xi without erosion of Xi and representing ∼20% of genes on the X, behaved largely as expected: of the escapee genes informative in all four lines, 14 of 18 were biallelic in every line (Tukiainen, Villani et al. 2017, Topa, Benoit-Pilven et al. 2024). However, a subset of what are believed to be constitutive escapees (e.g., USP9X and DDX3X) were biallelically expressed in GeDiX697 and GeDiX927 yet monoallelic in GeDiX395 and GeDiX002, a state supported by very high coverage (**Table S7**). Line-specific behavior was also evident at other escapees: PCDH19 was biallelic only in GeDiX002 and monoallelic in the other three lines, whereas TCEANC escaped in three of four lines, losing biallelic expression uniquely in GeDiX002. GeDiX002 thus both gained and lost escape at individual loci, underscoring the locus- and line-specific nature of these transitions. Conversely, KLHL4, classified as subject to X-inactivation in human reference tissues (Tukiainen, Villani et al. 2017, Topa, Benoit-Pilven et al. 2024), was robustly biallelically expressed in all four lines (**Table S6**).

The spatial distribution of escape, however, differed markedly between lines. Biallelic (escaping) genes were increasingly concentrated on the short arm (Xp) in the more eroded lines: the fraction of escaping genes on Xp rose from 25% in the most intact line (GeDiX697) to 37–39% in the more eroded lines (GeDiX002, GeDiX927), while escape on the long arm (Xq) declined over the same series (20% to 15%). A stable set of 13 genes escaped in all four lines (including 7 pseudoautosomal genes), whereas 12 genes gained biallelic expression specifically in the terminal line GeDiX927 (e.g., CLCN4, MID1, XIAP, PIGA), indicating locus-specific reactivation accompanying advanced erosion. We also examined specific loci known for either maintenance of monoallelic expression or susceptibility to erosion. For instance, *ATRX,* located approximately 4MB 3’ to XIST, is known to maintain monoallelic expression - in our data it remains entirely inactive on the Xi with only one read (in GeDiX697) from the alternate allele detected across all four female lines compared to 13,579 reads from the Xa (**Table S6**). This is in contrast to erosion-susceptible *USP9X* with Xi expression frequencies of 0.005, 0.007, 0.212, and 0.358 across GeDiX002, GeDiX395, GeDiX697, GeDiX927, respectively. Though unclear of its functional role in the process, the expression of XACT is a sensitive readout of the changing chromatin environment on the Xi (Vallot, Ouimette et al. 2015, Motosugi, Okada et al. 2021) – Xi expression frequencies for XACT were 0.037, 0.278, 0.309, and 0.487 for GeDiX lines 697, 395, 002, and 927, respectively, tracking with the order of erosion defined by the Barr body analysis.

Other key loci of interest that are dynamic, early events in erosion of Xi include the *DXZ4* macrosatellite repeat and the *FIRRE* locus as each is involved in the mechanism of XCI (Bansal, Kondaveeti et al. 2019). At *FIRRE* (**Figure 5E**), we identified 20 allelic DMRs across the locus, one of these was at the promoter of this Xa-expressed gene and was hypomethylated on the Xa. The remaining 19 DMRs were hypomethylated on the Xi, two upstream of the transcription start site and 17 within the gene body. The extent of Xi hypomethylation correlated with Barr body status, with GeDiX697 with the greatest extent of Xi hypomethylation and GeDiX927 with the least. The haplotype-resolved assembly of the *FIRRE* locus recovered the same local-repeat families previously described (Hacisuleyman, Shukla et al. 2016). Intersecting the repeats with the *FIRRE* DMRs, the most abundant repeat (R0) overlapped the majority of DMRs. There was no evidence of R0 copy number polymorphism across the GeDi lines, these were invariant across all haplotypes. The DMRs also coincided with CTCF binding sites: 17 of the 20 *FIRRE* DMRs overlapped a CTCF-bound regulatory element, predominantly distributed across the gene body (**Figure 5E**).

The DXZ4 repeat anchors the superloops structural involved in Xi and is in a euchromatic state, hypomethylated and bound by CTCF, on the Xi, whereas on the Xa these microsatellite repeats are in constitutive heterochromatin with hypermethylation (Chadwick 2008). Methylation across the *DXZ4* tandem-repeat array showed a periodic, wave-like pattern (**Figure 5F**). Because *DXZ4* is a polymorphic microsatellite that is collapsed within a 100 kb bridged gap in the hg38 reference, we quantified its copy number per haplotype directly from the phased assemblies (**Figure 5F**). Array size varied substantially across haplotypes, ranging from 28 to 84 monomer copies (84–249 kb). Reporting the copy-number values as observed: the Xa haplotype carried the greater number of copies in all four female lines (though in the most eroded line, GeDiX927, the two haplotypes were near-identical at 78 and 77 copies). The most eroded line, GeDiX927, also had the highest combined copy number across both haplotypes, the highest count in a single haplotype, and the smallest difference in copy number between its two haplotypes of the four lines.

## Discussion

The GeDi panel addresses a critical gap in the iPSC field by providing a well-characterized and quality-controlled resource of human iPSCs that captures a wide breadth of global human genetic variation with contributions from (in order of descending frequency) African, admixed American, East Asian, South Asian, Middle Eastern, and European ancestry - the 14,027,638 total unique SNPs across these eight lines are illustrative of this heterogeneity. Our systematic characterization across functional, genomic, transcriptomic, and epigenomic dimensions, including haplotype-resolved genome assemblies, demonstrates that the panel captures a sub-set of human genetic and phenotypic variation. Experience from model organism research has long shown that working across multiple genetic backgrounds is essential for distinguishing robust biological findings from background-specific artifacts — a principle that the iPSC field is only beginning to apply systematically (Pantazis, Yang et al. 2022).

### Quality-controlled heterogeneous iPSC lines for functional genomic studies

All eight GeDi lines and KOLF2.1J passed a comprehensive quality control pipeline including Sendai virus clearance, proliferative capacity, p53 DNA damage response, and XCI maintenance in female lines, and were successfully edited by CRISPR-Cas9. The intact p53 response is particularly important given the documented enrichment of dominant-negative TP53 mutations in iPSC repositories, which can compromise the validity of disease models (Merkle, Ghosh et al. 2017). Similarly, whole-genome sequencing identified a high-impact BCOR splice donor variant in GeDiY518.1J, consistent with the strong positive selection for BCOR mutations documented across large iPSC repositories during reprogramming (Rouhani, Zou et al. 2022). The isogenic corrected line GeDiY518.1.1J, generated here by prime editing, provides a clean comparison that unambiguously attributes the altered colony morphology, distinct transcriptomic state, and proliferative advantage of GeDiY518.1J to BCOR loss of function rather than donor background.

Proliferative capacity is a critical but often underappreciated variable when working with multiple lines in parallel. In pooled designs such as village cultures, differences in doubling time can bias readouts toward faster-growing donors, and growth advantages conferred by culture-adaptive mutations — as exemplified by the BCOR variant in GeDiY518.1J — can further skew representation, underscoring the importance of monitoring both proliferative capacity and genomic integrity in multi-line experiments. The doubling times we observed — ranging from 23.6 h (GeDiY518.1J, BCOR mutant) to 32.3 h (KOLF2.1J) — reflect stable line-intrinsic phenotypes that investigators should account for when designing experiments; notably, these rates were measured under vitronectin/StemFlex conditions, and each line, owing to its distinct genetic background, may respond differently to alternative substrates or media formulations.

The most direct measure of pluripotency is quantifying lineage differentiation and here we did this via transcriptomic analysis by RNA-seq across multiple lineages. Traditionally, this has been restricted to trilineage differentiation (*i.e.*, the lineages of the embryo proper) but here we show the multiple advantages of including the extra-embryonic lineages in this analysis. First, this provides an expanded differentiation potential of iPSC lines - the trophoblast lineage being the first lineage to form in development - and is a feature of human pluripotent stem cells not shared with conventionally cultured mouse counterparts as was evident in the first human derivations (Thomson, Itskovitz-Eldor et al. 1998, Reubinoff, Pera et al. 2000). Second, the measured transcriptome expanded significantly by including the extra-embryonic lineages – noting both the PrS and the ExM had significantly more differentially expressed genes identified, 3,595 and 3,223, respectively, compared to Endo/Meso/CBO (from 1,600 - 2,196). Third, this lineage provides the opportunity for functional evaluation of imprinted loci as many imprinted genes are specifically expressed in the extra-embryonic lineages. Finally, and with respect to human phenotypic variation, it is anticipated that such variation would be over-represented in the extra-embryonic lineages as the placenta is the most divergent organ across mammalian species – such phenotypic variation is exemplified in the case of the PSG locus described here.

### Genetic and epigenomic sources of variation across the GeDi panel

The genetic heterogeneity captured by the GeDi panel has direct practical implications for functional studies: research relying on a single genetic background will systematically underrepresent the full spectrum of human regulatory variation, potentially missing ancestry-specific effects on gene expression, splicing, and cellular phenotypes (Carcamo-Orive, Hoffman et al. 2017, Kilpinen, Goncalves et al. 2017). This is directly demonstrated by our multi-lineage transcriptomic data — within each of the five differentiated lineages, a substantial number of genes showed stable, donor-intrinsic variability in expression that would be invisible in studies using a single reference line, while replicates from the same donor clustered tightly, confirming that this variability reflects genetically and/or epigenetically determined differences (Kilpinen, Goncalves et al. 2017). This begins to position the GeDi panel as a platform for eQTL studies in differentiated cell types (DeBoever, Li et al. 2017, Cuomo, Seaton et al. 2020, Jerber, Seaton et al. 2021) and cross-ancestry analysis of disease-relevant gene regulation that will only be enhanced as we continue to build out this resource.

We provide a comprehensive analysis of the epigenomic profile of the GeDi lines utilizing global genomic methylation profiles which the long-read chemistry provides, in addition to haplotype-resolved monoallelic/biallelic expression to provide functional confirmation of the impact of epigenetic imprint variation. Global genomic methylation profiles position the GeDi lines close to other reference ESCs and iPSCs and distal to the donor cells they were derived from. We identify variation in sites of genomic imprinting, at some specific locations genomic imprints are highly stable across all lines (e.g., *PEG10* locus and the *H19, IGF2, CDKN1C* locus) whereas other sites of known genomic imprinting are less stable, varying across lines, and contributing to expression differences between lines in the regulating genes (e.g., *MEG3*, *DLK1*) associated with these loci. This underscores the importance of imprinting analysis as a component of iPSC quality control (Nazor, Altun et al. 2012, Bar, Schachter et al. 2017) and the value of incorporating long-read chemistry in this analysis.

Much remains unknown of the mechanism of control of human XCI (and how it erodes in iPSC cultures) - it is known to differ from the much better-defined mechanism in the mouse. There has been a tendency to favor male iPSC lines, in part, due to unknowns around Xi/e however, it is essential to have female iPSC lines represented in biomedical research. Thus, we have extensively quantitated changes associated with Xi/e, the extent of which varies across the four female GeDi lines. Specifically characterizing *XIST* expression, quantitating monoallelic expression across the X, and utilizing genomic methylation to define the patterns at key genomic loci functionally involved in the superloop structure on the Xi. One of these key loci, and differentially methylated, is the macrosatellite repeat DXZ4. The assembled GeDi genomes enabled characterization of the polymorphic nature of this repeat, which ranged in size from 28 to 84 monomer copies (84–249 kb). Intriguingly, the Xa haplotype carried the greater number of DXZ4 repeats in all four female lines and with the most eroded line, GeDiX927, containing two haplotypes that were near-identical in repeat size. Whether repeat number and/or differential in size across haplotype plays any choice Xi or its subsequent rate of erosion would require further analysis. Continuing to track all these features related to Xi/e over passage length and across a broader number of GeDiX lines may provide future mechanistic insight into this epigenetic phenomenon.

### Haplotype-resolved assemblies and structural variation

The haplotype-resolved assemblies provide the most comprehensive genomic characterization of an iPSC panel spanning multiple ancestry groups to date, cataloging SVs and novel regions of absence of heterozygosity (AOH) that were previously inaccessible to short-read sequencing. This approach is consistent with the value of custom reference assemblies recently demonstrated for KOLF2.1J, where long-read-based assembly revealed SVs — including those overlapping neurodevelopmentally relevant genes — that were not detected by SNP array or short-read sequencing (Jerez, Rhie et al. 2026). Together, these findings underscore the importance of genome assemblies for characterization of reference iPSC lines. Within this framework, the PSG locus on chromosome 19 — an established region of population-level SV encoding glycoproteins expressed exclusively in syncytiotrophoblast and characterized by rapid evolution in primates with hemochorial placentation (Zimmermann and Kammerer 2021) — emerges as a hotspot of recurrent structural variation across the panel. Although long-read sequencing and haplotype-resolved assembly approaches have become increasingly routine, the majority of lines that have been sequenced are lymphoblastoid cell lines (Logsdon, Ebert et al. 2025), with limited potential for examination of biological questions. As iPSCs, the GeDi lines will provide the cellular resource in which to functionally address genomic variation revealed by their respective high-quality phased reference genomes, enabling differentiation to whatever cell type is anticipated to express the associated phenotype.

### Common Functional Variation Across Lines

In addition to raw and processed data, we provide access to a genome viewer (wge.jax.org) that can quickly identify lines containing SNPs of interest to the community, and assist in line selection for downstream investigations. It also serves as a useful tool to identify CRISPR guides. Some examples of functional SNPs represented in the GeDi lines include the common variant (rs1047303) known to alter the half-life of the steroidogenic enzyme HSD3B1 (Chang, Li et al. 2013) and a variant (rs4769613) upstream of *FLT1* that is the strongest genetic association with preeclampsia (McGinnis, Steinthorsdottir et al. 2017). Other SNPs represented in the GeDi lines, include the Type 2 Diabetes risk allele rs6444081 associated with reduced activity of a stress-responsive cis-regulatory element (Sokolowski, Kursawe et al. 2024) and rs9923231, a well-known functional *VKORC1* promoter polymorphism affecting warfarin sensitivity known to be enriched in East Asian populations (Ross, Bigham et al. 2010). These represent only a handful of examples of common variants represented and variable across the eight GeDi lines.

### Limitations and future directions

Limitations of the current study should be noted. The panel captures a wide breadth of global human genetic variation, however this limits statistical power for population-level association analyses within individual lineages. Expanding the panel to include additional donors that capture additional, here-to unsampled genetic diversity, particularly those from underrepresented populations (e.g., sub-Saharan African, Middle Eastern, and Indigenous American groups) would increase resolution, and is currently being actively pursued. It should also be noted that the GeDi panel is not intended to replace single reference line studies but to complement them — providing genetic context and population-level resolution that cannot be achieved with a single donor background while remaining fully compatible with existing workflows built around KOLF2.1J and other established reference lines. The differentiation protocols used here produce relatively early-stage cell types, and further maturation will be needed for disease modelling applications requiring more mature cellular identities. Finally, all functional characterization was performed under a single culture condition — vitronectin coating with StemFlex medium — and the response of individual lines to alternative substrates or media formulations remains to be determined.

An important future direction will be to systematically correlate genetic background with functional readouts across the panel. We have begun to establish these connections — for example, structural variants at the PSG locus correlate directly with changes in PSG gene expression, demonstrating that haplotype-resolved assemblies can identify genotype-to-expression relationships at complex loci. However, the downstream phenotypic consequences of such variants remain to be determined. More broadly, epigenetic instabilities such as XCI erosion or loss of imprinting — which vary across the panel — may contribute to inter-line differences in differentiation studies and could be misattributed to genetic effects if not accounted for. The integration of haplotype-resolved *XIST* promoter and *FIRRE*/*DXZ4* loci genomic methylation, allele specific expression, and H3K27me3 immunostaining provides a multi-modal approach to XCI assessment, demonstrating concordance between molecular and functional readouts and offering a framework for monitoring epigenetic stability across passages and experimental contexts. The GeDi panel, with its combination of haplotype-resolved genome assemblies, whole-genome methylation data, and multi-lineage transcriptomics, provides an ideal substrate for the integrative multi-omic analyses needed to disentangle these contributions.

Despite these limitations, the GeDi panel represents a meaningful advance in the availability of quality-controlled iPSC resources spanning multiple ancestry groups. By combining rigorous characterization, open distribution of cell lines and data, and compatibility with standardized genome editing workflows, the panel is designed to support functional genomic studies across human populations, broaden our understanding of human biology and disease, and bring enhanced rigor and reproducibility to functional genomic studies leveraging human iPSC-derived models applicable to human health and disease.

## Acknowledgements

We are first and foremost thankful to all donors who participated in this study and their willingness to make the derived products from their donations widely available to the research community. We gratefully acknowledge the JAX Genome Technologies services, the Cyberinfrastructure high-performance computing resources, and the JAX Flow Cytometry facility at The Jackson Laboratory; these shared services are supported in part by the JAX Cancer Center (P30 CA034196). We thank Nélio A. J. Oliveira for his contribution to the Sendai virus clearance flow cytometry assays. We thank Mallory Ryan and Kaitlyn Stewart for the SNP array analysis. We thank the UConn Health Clinical Research Center and Heather Vargas for coordinating donor recruitment and blood collection. We also thank Stefan Pinter for discussion on Xi erosion.

## Data Availability

The GeDi iPSC lines described in this study will be made publicly available through The Jackson Laboratory repository (https://www.jax.org/ipsc). Prior to public distribution, lines are available upon request to the corresponding author. Data have been deposited at the European Genome-phenome Archive (EGA), which is hosted by the EBI and the CRG, under accession number EGAS00001008380. Additional data are available upon request to the corresponding authors.

## Material and Methods

### Materials availability

- All GeDi iPSC lines generated in this study are available upon request prior to their public distribution through The Jackson Laboratory repository (https://www.jax.org/ipsc). Requests for materials should be directed to the corresponding authors.
- KOLF2.1J (product code: JIPSC001000), KOLF2.2J (product code: JIPSC003902) and their derivatives are available from The Jackson Laboratories at https://www.jax.org/ipsc.

### Donor recruitment and ethics

Healthy volunteer donors were recruited through the UConn Health Clinical Research Center under IRB protocol #24-023J-2. The study aimed to recruit donors representing diverse population groups based on self-reported race, ancestry, and ethnicity. Eligibility criteria included age 18–35 years, no history of malignancy, and willingness to consent to de-identified broad genomic data sharing and of derived iPSC lines including genomic sequencing data for general research use. The relatively young upper age bound was chosen to limit the possibility of capturing somatic genome variation that is known to increase with age (Bernstein, Spencer Chapman et al. 2024). Pregnant individuals were excluded. All participants provided written informed consent prior to enrolment. Donors received $50 compensation for their participation. Race and ethnicity information were collected using the categories established under the 1997 OMB Revisions to the Standards for the Classification of Federal Data on Race and Ethnicity. Donors were also given the option to provide additional detail on self-reported ancestry by selecting from specific global geographic regions. In some instances, the country of origin for the donor family was provided by the donor. Whole blood donor samples were de-identified prior to initiation of reprogramming.

### iPSC reprogramming from PBMCs

Each donor attended a single study visit at the UConn Health Clinical Research Center during which a blood sample was collected by standard venipuncture. Peripheral blood mononuclear cells (PBMCs) from healthy donors representing diverse population groups were isolated from whole blood by density gradient centrifugation using SepMate-15 tubes (STEMCELL Technologies, Cat# 86415) loaded with Ficoll-Paque (4 mL per tube), (Chou, Mali et al. 2011). Blood was diluted 1:1 with PBS and layered onto the Ficoll before centrifugation at 1,200 × g for 10 minutes. The mononuclear cell layer was collected and washed twice in PBS + 2% FBS. PBMCs were enumerated by haemocytometer and either cryopreserved in Knockout Serum Replacement (KOSR; Thermo Fisher Scientific, Cat# 10828-028) + 10% DMSO or used directly for EPC expansion.

For EPC expansion, cryopreserved PBMCs were thawed by adding 0.5 mL thawed cells to 2 mL pre-warmed PBS + 2% FBS and centrifuged at 300 × g for 5 minutes. After a second wash in PBS + 2% FBS, cells were resuspended in StemSpan SFEM II (STEMCELL Technologies, Cat# 09605) supplemented with the following recombinant human cytokines at the indicated final concentrations: SCF (C-Kit Ligand; 100 ng/mL; Thermo Fisher Scientific, Cat# PHC2111), FLT-3 Ligand (100 ng/mL; Cat# PHC9414), IL-3 (20 ng/mL; Cat# PHC0034), and IL-6 (20 ng/mL; Cat# PHC0065), together with Primocin (1:500, 20 µL per 10 mL; InvivoGen, Cat# ant-pm-1) and Penicillin/Streptomycin (1:100; Life Technologies, Cat# 15140-122). Cells were plated in two untreated wells of a 24-well plate and medium was changed every 2 days. Primocin was removed on day 6. Cells were cultured for 9 days total (D−9 to D0) to enrich for EPCs.

On day 0 of transduction, 50,000 EPCs per well were transduced with the CytoTune-iPS 2.0 Sendai Reprogramming Kit (Thermo Fisher Scientific, Cat# A1378001) using the three reprogramming vectors — CytoTune 2.0 KOS (encoding KLF4, OCT3/4, and SOX2), CytoTune 2.0 hc-Myc, and CytoTune 2.0 hKLF4 — at MOIs of 5, 5, and 3 respectively. Viral volumes were calculated per the manufacturer’s formula:

Volume of virus (µL) = MOI (CIU/cell) × number of cells / titer (CIU/mL) × 10 ³ (mL/µL)

Virus was added to cells in StemSpan SFEM II + Penicillin/Streptomycin and transduction efficiency was enhanced by spinoculation at 2,250 rpm for 90 minutes at room temperature. On day 1, virus was removed by centrifuging cells at 300 × g for 10 minutes, aspirating the supernatant, and resuspending in 1 mL fresh StemSpan SFEM II + Penicillin/Streptomycin per well.

On day 3, transduced cells were collected, centrifuged at 300 × g for 10 minutes, resuspended in ReproTeSR medium (STEMCELL Technologies, Cat# 05926) supplemented with the above cytokines (SCF, FLT-3 Ligand, IL-3, IL-6) and CloneR2 (STEMCELL Technologies), and replated on Matrigel-coated wells (Corning ES-qualified Matrigel, Cat# 354230). Medium was changed every 2 days. From day 7–12, when small iPSC colonies became visible, medium was transitioned to mTeSR Plus (STEMCELL Technologies, Cat# 100-0276) supplemented with CloneR2, and changed daily. Once colonies were well established, medium was transitioned to StemFlex (Thermo Fisher Scientific, Cat# A3349401) supplemented with RevitaCell (REV; 1X; Thermo Fisher Scientific), which was maintained for all subsequent passages.

At approximately day 14, wells were passaged using Accutase (10 minutes at 37°C) and replated on Matrigel-coated wells in mTeSR Plus + CloneR2. Medium was changed daily. Upon confluence, cells were passaged at 1:10 and two vials were cryopreserved at passage 1 in KOSR + 10% DMSO. Cells were further expanded and master stock vials cryopreserved at passage 6. Single-cell subcloning was performed at passage 5, and lead clonal sublines were selected based on SNP array comparison to the parental PBMC sample to confirm genomic integrity.

### iPSC maintenance and cell culture

All iPSC lines were maintained under feeder-free conditions on vitronectin-coated plates (Vitronectin XF, STEMCELL Technologies, Cat# 07180) in StemFlex medium (Thermo Fisher Scientific, Cat# A3349401) at 37°C, 5% CO. Plates were coated by diluting vitronectin 1:100 in PBS and incubating for at least 1 hour at room temperature before use; coated plates were stored dry at 4°C for up to 3 months. Cells were passaged every 3–4 days using ReLeSR (STEMCELL Technologies, Cat# 05872) when colonies reached 70–80% confluency. Briefly, spent medium was aspirated and cells were washed once with room-temperature DPBS (Thermo Fisher Scientific, Cat# 14190-094). ReLeSR (2 mL per well of a 6-well plate) was added and incubated for 5 minutes at room temperature, then aspirated. StemFlex medium (3 mL) was added, and cells were gently detached using a cell lifter (Corning, Cat# 3008) and broken into small clumps by pipetting 2–3 times. Cells were passaged at a 1:15 dilution onto fresh vitronectin-coated plates. Medium was changed every 2 days until confluence, typically within 3–4 days.

When single-cell suspensions were required — for nucleofection, single-cell cloning, or seeding of IncuCyte proliferation assays — cells were dissociated with Accutase (STEMCELL Technologies) for 7–10 minutes at 37°C, and StemFlex medium supplemented with RevitaCell (1X; Thermo Fisher Scientific) or CloneR™2 (1X; STEMCELL Technologies #100-0691) was used for replating to support survival of single cells.

### Sendai virus clearance assay

Residual Sendai virus was assessed by intracellular flow cytometry using a intracellular antigen staining protocol. Cells were dissociated to single-cell suspension using Accutase (7–10 minutes at 37°C), washed in PBS + 2% FBS + 2 mM EDTA (FACS buffer), and transferred to a V-bottom 96-well plate (200 μL per well). Cells were stained with LIVE/DEAD Fixable Aqua Dead Cell Stain (1:1000; Thermo Fisher Scientific, Cat# L34957) for 10–15 minutes in the dark, then washed with FACS buffer. Cells were fixed and permeabilized using the BD Cytofix/Cytoperm Kit (BD Biosciences, Cat# 554714) according to the manufacturer’s instructions. Following Fc blocking (Human Fc Block, BD Biosciences, Cat# 564765), cells were stained with an anti-Sendai virus HN antibody conjugated to AF647, diluted in BD Perm/Wash buffer, for 30 minutes on ice. Cells were washed twice and resuspended in FACS buffer for acquisition. An early-passage iPSC pool at passage 3 post-reprogramming served as a positive control (69.2% HN-AF647+). Data were acquired on a BD FACSymphony A5 cell analyzer (BD Biosciences) and analyzed using FlowJo^TM^10. The percentage of HN-AF647+ cells was calculated within the live singlet gate.

### IncuCyte live-cell proliferation assay

iPSC lines were seeded at 2,500 cells per well in vitronectin-coated 96-well plates in StemFlex + CloneR™2. Plates were transferred to an IncuCyte S3 live-cell imaging system (Sartorius) and imaged every 6 hours over 84 hours using a 4X objective. Phase-object confluency (%) was quantified using AI-based cell segmentation in the IncuCyte analysis software. Three independent experiments were performed for each line. Growth curves were normalised to the 12-hour timepoint to account for initial seeding variation. Doubling times were calculated by fitting a log-linear model to the exponential growth phase (12–60h) in R (v2024.12.1). Statistical comparisons of doubling times between lines were performed by one-way ANOVA with Tukey post-hoc correction using the rstatix package.

### p53 DNA damage response assay

To assess p53 pathway integrity, a stable mCherry reporter line was generated by piggyBac transposition. The reporter construct PG13-mCherry (Addgene, #195291), containing 13 tandem p53 consensus binding sites (p53 response elements) upstream of an mCherry open reading frame, was co-delivered with the hyperactive piggyBac transposase plasmid pCMVhyPBase into iPSCs by nucleofection (Lonza 4D Nucleofector, program CA-137). Briefly, iPSCs were dissociated with Accutase, and 2.2 × 10 cells per reaction were resuspended in 20 μL P3 Primary Cell Nucleofector Solution (Lonza). The nucleofection mixture contained 0.375μg pCMVhyPBase and 0.625μg PG13-mCherry per reaction. Following nucleofection, cells were plated on vitronectin-coated 12-well plates in StemFlex + CloneR2. Stable reporter-positive cells were selected with puromycin (Sigma-Aldrich) for 7 days at a concentration of 0.5 µg/mL, then expanded and used for downstream assays.

For doxorubicin treatment, iPSCs were seeded on vitronectin-coated plates and cultured for 24 hours prior to treatment. Doxorubicin (Sigma-Aldrich) was added at a final concentration of 50 nM and cells were incubated for 24 hours at 37°C. Cells were then dissociated with Accutase, washed in DPBS and resuspend in PBS + 2% FBS + 2 mM EDTA (FACS buffer). mCherry fluorescence was measured by flow cytometry. A TP53 KO iPSC line (KOLF_BFP_TP53KO, generate in the Skarnes Lab, unpublished) served as a negative control for p53-dependent reporter induction. The percentage of mCherry+ cells was calculated within the singlet gate using FlowJo^TM^10.

### X chromosome inactivation — H3K27me3 immunofluorescence

XCI status was quantified by H3K27me3 immunofluorescence. iPSC lines were seeded at approximately 30,000 cells per well in vitronectin-coated 96-well PhenoPlate plates (Fisher Scientific, Cat# 50-209-9836) and grown to ∼75% confluency. Cells were fixed with 100 μL of 4% paraformaldehyde (PFA) in PBS for 10 minutes at room temperature, washed once with PBS, and stored in PBS at 4°C until staining. For permeabilization and blocking, cells were incubated with 100 μL of 0.5% Triton X-100 in 4% BSA for 30 minutes at room temperature. After three washes with PBS, cells were incubated overnight at 4°C with anti-H3K27me3 primary antibody (Cell Signaling Technology, clone C36B11, 1:800 in 4% BSA). Following three PBS washes, cells were incubated with Alexa Fluor 488-conjugated secondary antibody (1:500 in 4% BSA) and Hoechst 33342 for 60 minutes at room temperature. After three final washes, plates were imaged on the Opera Phenix high-content imaging system (PerkinElmer) at 40X objectives. Automated image analysis was performed using a custom pipeline: nuclei were segmented based on Hoechst signal, and H3K27me3 foci were identified within each nucleus using spot detection. XCI+ nuclei were defined as nuclei containing at least one discrete H3K27me3 focus, and the percentage of XCI+ nuclei was calculated as (XCI+ nuclei / total selected nuclei) × 100.

### GCM1 CRISPR-Cas9 knock-out — PTC+2 NGGV strategy

GCM1 knock-out was introduced across all GeDi lines and KOLF2.1J using the PTC+2 (premature termination codon + 2 frameshift) strategy with an NGGV donor design, as described in the JAX DPC protocol (Pellegrino, McDonough et al. 2025), and adapted from Skarnes et al. (Skarnes, Pellegrino et al. 2019) and the MorPhiC consortium (Adli, Przybyla et al. 2025). A guide RNA targeting an early constitutive exon of GCM1 (at least 35 amino acids downstream of the start codon, common to all transcript isoforms) was designed to minimize off-target activity and selected for the sense strand:

GCM1_NGGV_sg1: 5’-CCGCGACTGTCTCGCAGAGG-3’ (PAM: AGG)

A 100-mer asymmetric antisense ssODN donor (GCM1_NGGV_A_t) was designed to introduce a TGA stop codon followed by two degenerate nucleotides in an NGGV configuration immediately downstream, resulting in a net +2 frameshift and preventing creation of a novel splice donor site. A degenerate base B (representing C/G/T; not A) was incorporated at a defined position within the edit to allow unambiguous discrimination of homozygous edited clones from wild-type by Sanger sequencing, as homozygous clones will lack the A base present in the wild-type sequence at that position:

5’-GCTGCTGCTTCTGCCGGGCCTTGTCACAGATGGCAGGTCTCAGGTAGATCTTGCGCC BCCNTCTCATGCGAGACAGTCGCGGCCGCACACCACCACACCC-3’

Cas9 RNP complexes were assembled by incubating recombinant SpCas9 protein with chemically modified synthetic sgRNA for 5–10 minutes at room temperature. For nucleofection, iPSCs were dissociated with Accutase and 2.2 × 10 cells per reaction were resuspended in 20 µL P3 Primary Cell Nucleofector Solution with supplement (Lonza, ratio 4.5:1 basal to supplement). RNP complex and ssODN donor were added to the cell suspension and transferred to a 16-well Nucleocuvette Strip (Lonza). Nucleofection was performed using the Lonza 4D Nucleofector with program CA-137. Immediately following nucleofection, cells were transferred to vitronectin-coated plates in StemFlex + CloneR2 and Alt-R HDR Enhancer V2 (IDT). Cells were incubated at 32°C for 3 days before transfer to 37°C. At 24h post-nucleofection, and every other day thereafter, the media was replaced with only StemFlex. Upon reaching near confluency, cells were single-cell-plated into Synthemax-coated 10cm dishes as described above. At Day 10, 24 colonies per cell sub-line were manually picked as previously described (Skarnes, Pellegrino et al. 2019) and incubated in Vitronectin-coated 96-well plates for 4–5 days before being frozen down. Clones were genotyped by Sanger sequencing using primers flanking the target site (outside the ssODN sequence). Genotypes were classified as homozygous PTC+2 (HOM), hemizygous (Hemi), heterozygous (Het), NHEJ indel, or wild-type (WT) based on sequence trace deconvolution.

### BCOR variant correction by CRISPR prime editing

The BCOR splice donor variant (chrX:40054254, A>G) identified in GeDiY518.1J was corrected to the reference allele (G>A) by CRISPR prime editing. An engineered pegRNA (epegRNA) was designed manually following published design guidelines, consisting of a 22-nucleotide spacer (5’-TTCTATGGCAGCTCTGTTTGTG-3’) adjacent to an NGG PAM site targeting BCOR exon 13, a reverse transcriptase template (RTT) of 12 nucleotides (5’-TCGAATGGTGTT-3’) encoding the reference A allele, and a primer binding site (PBS) of 13 nucleotides (5’-TGTCTCGACGGTA-3’). The tevoPreQ1 motif was appended to the 3’ end of the epegRNA to improve stability. The epegRNA was custom-synthesised with chemical modifications by Integrated DNA Technologies (IDT). PEmax and MLH1dn were custom-synthesised as in vitro-transcribed mRNA by TriLink Biotechnologies, based on the sequences in Addgene plasmid #174820, using unmodified bases with CleanCap technology.

For nucleofection, GeDiY518.1J cells were dissociated with Accutase (STEMCELL Technologies, Cat# 07920) and centrifuged at 300 × g for 5 minutes in StemFlex + CloneR2 (STEMCELL Technologies, Cat# 100-0691). After counting by haemocytometer, 2 × 10 cells were transferred to a V-bottom plate, centrifuged at 300 × g for 5 minutes, and the medium aspirated. The cell pellet was resuspended in 20 µL P3 Primary Cell Nucleofector Solution (Lonza, ratio 4.5:1 basal to supplement) containing PEmax mRNA and epegRNA. The suspension was transferred to a 16-well Nucleocuvette Strip and nucleofected using a Lonza 4D Nucleofector X Unit (Cat# AAF-1003X) with program CA-137. Immediately following nucleofection, cells were replated onto vitronectin-coated 24-well plates in StemFlex + CloneR2 (STEMCELL Technologies, Cat# 100-0691).

Following recovery, single-cell clones were isolated and screened by Sanger sequencing at the target locus. The corrected line (GeDiY518.1.1J) was confirmed to carry the reference A allele, expanded, and used for transcriptomic and morphological comparisons with the parental BCOR mutant line GeDiY518.1J.

### Illumina short-read whole-genome sequencing and variant calling

Genomic DNA samples were submitted to the JAX Genome Technologies service for Illumina short-read whole-genome sequencing (WGS). Briefly, 200 ng of DNA was sheared to 400 bp using a Covaris R230 focused-ultrasonicator and size-selected using AMPure beads (Beckman Coulter). Fragments were end-repaired, A-tailed, and ligated to Illumina Unique Dual Adapters using the KAPA Hyper Prep Kit (Roche) followed by 5 cycles of PCR amplification. Sequencing was performed on an Illumina NovaSeq X Plus platform generating paired-end reads of 2 × 151 bp.

Raw sequencing reads (FASTQ files) were processed through a standard GATK-based variant calling pipeline: reads were aligned to the GRCh38 reference genome and BAM and VCF files were generated following GATK best practices (pipeline details available at https://github.com/TheJacksonLaboratory/jds-nf-workflows/wiki/WGS-Pipeline-ReadMe).

Resulting variant calls were loaded into the WGE genome browser application (https://wge.jax.org) for interactive visualisation of single nucleotide variants across selected lines. BAM files were used for region-specific analysis using the Broad Institute Integrative Genomics Viewer (IGV; https://igv.org/app/).

### Emedgene analysis

FASTQ files were aligned to human reference genome hg38 (Illumina reference file: hg38-alt_masked.graph.cnv.hla.rna_v4.tar.gz). Variant calling was performed using the Illumina DRAGEN Germline Whole Genome v4.3.13-1 pipeline within the Illumina Connected Analytics (ICA) platform. Variant annotation and tertiary analysis were performed within the Illumina Emedgene platform (v38.3.7). All variants were interpreted according to modified ACMG/AMP guidelines (PMID: 25741868, 40568962, 38814327). Variants classified as pathogenic or likely pathogenic were reported across all genes, while variants of uncertain significance (VUS) were reported only for a pre-specified gene list of important iPSC pluripotency markers: BCOR, TP53, NANOG, DNMT3B, GBX2, NANOGP8, SP8, PEG3, ZIC1, POU5F1, SOX2, KLF4, MYC, MKI67, PCNA, MCM2, FSTL1, TOP3B, FGF4, FGD4, FGF2, TACO1, and FGFR1–4.

### Ancestry and genetic diversity of the donor iPSC panel

#### Population-descriptor terminology

Donor race, ethnicity, and ancestry were self-reported by each donor and are presented as such (**Table S8**). Genetic results are described in terms of genetic similarity to reference groups of the 1000 Genomes (1KG) Project, rather than as intrinsic or continental ancestry, following the framework of the National Academies of Sciences, Engineering, and Medicine (2023).

Candidate lines were selected using donor self-reported ancestry; genetic data were used subsequently to evaluate the diversity of the resulting panel rather than to assign ancestry to individual lines. Reference samples retain their original 1KG population codes (e.g. ASW, MXL) throughout. Primary labels are aggregated into the five secondary labels defined by the 1000 Genomes Project (AFR, African; AMR, American; EAS, East Asian; EUR, European; SAS, South Asian) only where an aggregate analysis is required, specifically for visualizing reference samples in PCA space and for summarizing genetic diversity metrics (**Figures 1D**, **1E**); ancestry proportions are estimated and reported at the level of the 26 primary labels.

For conciseness we use the abbreviation-plus-“-like” convention recommended in the NASEM report: 1KG-AFR-like, for example, denotes genetic similarity to the 1KG African reference group. These codes are used as genetic-similarity references, not as bounded continental categories.

#### SNP-array genotyping and data processing

Genomic DNA was extracted from all GeDi iPSC lines and parental PBMC samples using the Takara NucleoSpin Tissue kit, eluting in 50 µL BE buffer. DNA concentration was quantified using the Qubit Fluorometer (Thermo Fisher Scientific), with a minimum quantity threshold of 50 ng/µL for array submission. SNP array genotyping was performed using the Illumina GDA Cyto-8 array at the JAX Advanced Precision Medicine Lab (APML, The Jackson Laboratory, USA), producing genotype call (.gtc) files oriented to the GRCh37 reference assembly. BioNano VIA software was applied to screen array data quality, detect copy number variants (CNVs) and regions of absence of heterozygosity (AOH), and provide a visualization and annotation platform. All CNV calls were manually reviewed, with particular attention to the 20q region, sex chromosomes, and AOH regions. For clonal subline selection, parental PBMC samples and all corresponding clonal sublines were co-analysed in VIA to identify de novo CNVs arising during reprogramming or subcloning. Sublines were selected for the GeDi panel on the basis of genomic concordance with the parental PBMC sample, absence of large chromosomal gains or losses, and absence of novel AOH regions. Lead sublines were advanced to downstream characterization only upon passing all QC thresholds.

For downstream genomic analysis, GTC files were converted to VCF using the Illumina Array Analysis CLI v2.1.0 with the BPM and CSV manifests and the GRCh37 reference FASTA. Because microarray probes can interrogate either genomic strand, variants were restricted to the sense (plus) strand prior to analysis. Each per-chromosome VCF was intersected with the Ensembl GRCh37 gene annotation (Homo_sapiens.GRCh37.87) using bedtools v2.31.1 (Quinlan and Hall 2010), retaining only plus-strand variants. Filtered variants were left-aligned and normalized against the hs37d5 reference assembly, with multiallelic sites decomposed and duplicate records removed. Normalized calls were converted to PLINK binary format (PLINK v1.9), (Purcell, Neale et al. 2007) with allele order preserved. Only autosomes (chr1–22) were carried forward; sex chromosomes were excluded.

#### Individual-specific identity markers

To enable verification of line identity and detection of sample mix-ups in downstream experiments, we derived a panel of private single-nucleotide variants (“genetic fingerprints”) for each line from the donor blood SNP-array genotypes. From the merged multi-sample array call set (GRCh37/hg19), cohort singletons were defined as sites at which the alternate allele occurred exactly once across the cohort (INFO/AC = 1), and each singleton was assigned to its carrier— the single individual with a non-reference, non-missing genotype—using bcftools v1.12. Because these genotypes are array-derived, candidate markers were restricted to high-confidence heterozygous calls by retaining only sites with a B-allele frequency between 0.45 and 0.55 (FORMAT/BAF), removing intensity-imbalanced or copy-number-affected probes.

For the eight GeDi donor lines, each candidate marker was then validated as truly private by querying, at the marker position, the blood- and iPSC-derived genotype call sets of every other individual in the cohort; any marker whose genotype was also observed in another individual was discarded. The retained private variants were lifted from GRCh37 to GRCh38 (CrossMap, hg19-to-hg38 chain), re-normalized against the GRCh38 reference, and annotated against GENCODE gene features (bedtools intersect. Two marker sets were generated per line: a “genomic” set comprising all private variants, and a “transcriptomic” set restricted to variants in transcribed exonic regions (CDS, UTR or exon), i.e. those potentially detectable in RNA-based assays.

#### Reference panel and principal component analysis

Reference genotypes were obtained from the 1000 Genomes Project phase 3 (release 20130502, GRCh37; phased SHAPEIT2/MVNcall integrated callset), (Genomes Project, Auton et al. 2015) and processed through the identical normalization and PLINK-conversion workflow described above. The donor and reference datasets were then reduced to their shared set of variants and merged. Principal component analysis was carried out on the merged dataset using PLINK. Donor lines were visualized in the space of the 1000 Genomes reference and labeled by the five 1000 Genomes reference groups (AFR, AMR, EAS, EUR, SAS).

#### Supervised admixture analysis

Per-line proportions of genetic similarity to each reference population were estimated with ADMIXTURE 1.3.0 in supervised mode on the merged PLINK dataset. The number of components was fixed at K = 26 to match the number of reference populations, so that each donor genome was modeled as a mixture of similarity to the 26 reference populations (Genomes Project, Auton et al. 2015). Per-line proportions are shown as stacked bars at the population level.

#### Genetic diversity from whole-genome sequencing

Whole-genome germline variants for each line were called with the Illumina DRAGEN pipeline v4.3.6. For each per-sample hard-filtered VCF, PASS variants were left-aligned and decomposed into biallelic records against the reference genome. The nine per-sample VCFs were merged into a multi-sample callset, and allele counts and frequencies (AC, AN, AF) were recomputed with bcftools v1.12 (Li 2011).

Per-line summaries—counts of heterozygous, homozygous-alternate and homozygous-reference genotypes, the heterozygosity rate (heterozygous genotypes / total genotyped sites), and the heterozygous-to-homozygous-alternate ratio—were computed in Python with scikit-allel (Chen and Huang 2026). The total number of non-reference variants per line was tabulated with bcftools v1.12, and private variants were defined as cohort-singletons (sites at which exactly one line carried the alternate allele, INFO/AC = 1). Per-sample heterozygosity was additionally estimated with VCFtools v0.1.16 (Danecek, Auton et al. 2011). For visualization, three per-line metrics—total variants, heterozygosity rate and private-variant count—were plotted and lines were grouped by donor self-reported ancestry (Africa, Americas, East Asia, Europe, Middle East, South Asia).

#### Directed differentiation of GeDi iPSC lines

To assess differentiation capacity and overall reprogramming quality, each line was maintained in the undifferentiated state (Und) and directed toward five lineages spanning embryonic and extra-embryonic fates: definitive endoderm (Endo), mesoderm (Meso), dorsal-forebrain cortical organoid (CBO), extra-embryonic mesenchyme (ExM) and primitive syncytium (PrS).

All differentiation protocols were performed on KOLF2.2J and GeDi lines maintained on Matrigel-coated plates in mTeSR Plus medium (STEMCELL Technologies, Cat# 100-0276) with daily medium changes until approximately 70–80% confluency at day 0. For each protocol, cells were dissociated to single-cell suspension using Accutase, counted by haemocytometer using trypan blue exclusion, and replated onto fresh Matrigel-coated plates in mTeSR Plus supplemented with Y-27632 ROCK inhibitor (10 µM) at the appropriate seeding density for each lineage. Material for RNA sequencing was collected at the endpoint of each protocol: undifferentiated iPSCs in the pluripotent state; definitive endoderm and mesoderm after 5 days of monolayer induction; primitive syncytium and extra-embryonic mesenchyme after 6 days of trophoblast-directed differentiation; and dorsal-forebrain cortical organoids after 9 days.

Each line was profiled in biological triplicate for every condition. Two lines (GeDiX697 and GeDiY634) failed to yield ExM material and were excluded from that lineage, and two individual replicates were removed as outliers during quality control (one Endo replicate of GeDiY016 and one Meso replicate of GeDiX395). After these exclusions, 154 RNA-seq libraries were retained for analysis (Und, n = 27; Endo, n = 26; Meso, n = 26; CBO, n = 27; ExM, n = 21; PrS, n = 27). All protocols are deposited on protocols.io and detailed below.

#### Mesoderm induction

Mesoderm differentiation was performed as described in the JAX DPC protocol (C., A.J. et al. 2025). Briefly, iPSCs were seeded in mTeSR Plus medium onto Matrigel-coated 6-well plates and differentiated towards mesoderm over 5 days using STEMdiff^TM^ Mesoderm Induction Medium (STEMCELL Technologies, Cat# 05221). Medium was changed daily throughout the differentiation. At the endpoint, cells were harvested for RNA extraction and downstream transcriptomic analysis.

#### Definitive endoderm induction

Definitive endoderm differentiation was performed as described in the JAX DPC protocol (C., A.J. et al. 2025). Briefly, iPSCs were seeded in mTeSR Plus onto Matrigel-coated 6-well plates and differentiated over 5 days towards definitive endoderm using STEMdiff^TM^ Definitive Endoderm Kit (STEMCELL Technologies, Cat# 05110) with medium changed daily. At the endpoint, cells were harvested for RNA extraction and downstream transcriptomic analysis.

#### Primitive syncytium differentiation

Primitive syncytium differentiation was performed as described in the JAX DPC protocol (Liu, Tan et al. 2025). This protocol differentiates iPSCs towards primitive syncytium, a cell type within the trophoblast lineage, by seeding the appropriate density onto Synthemax-coated 6-well plates in StemFlex Medium and culturing cells with TE differentiation medium supplemented with BMP4 (100 ng/mL), SU5402 (20 uM), and A-83-01 (1 uM) for 6 days. At the endpoint, cells were harvested for RNA extraction and downstream transcriptomic analysis.

#### Extra-embryonic mesenchyme differentiation

Extra-embryonic mesenchyme differentiation was performed as described in the JAX DPC protocol (Z., J. et al. 2025). This protocol differentiates iPSCs towards extra-embryonic mesenchyme, a cell type within the trophoblast lineage, through seeding the appropriate density onto Synthemax-coated 6-well plates in StemFlex Medium and culturing cells with TE differentiation medium supplemented with BMP4 (100 ng/mL), SU5402 (20 uM), and Activin A (100 ng/mL) for 6 days. At the endpoint, cells were harvested for RNA extraction and downstream transcriptomic analysis.

#### Dorsal forebrain organoids

Dorsal forebrain organoids were generated as described in the JAX DPC protocol (A., C. et al. 2025). Briefly, iPSCs were maintained on Matrigel-coated plates and passaged using dispase, with spontaneously differentiated cells removed prior to organoid induction. Embryoid bodies were formed in suspension using an orbital shaker platform and organoids were guided through a dorsal forebrain patterning protocol using dual SMAD inhibition and WNT inhibition to generate cortical organoids. Organoids were maintained in various neural differentiation media for 9 days until harvest for RNA extraction and downstream transcriptomic analysis.

#### RNA extraction, library preparation and sequencing

Total RNA was extracted at each differentiation endpoint using the RNeasy Plus Mini Kit (Qiagen) for embryonic lineages and the PicoPure RNA Isolation Kit (Applied Biosystems) for extra-embryonic lineages. Strand-specific, rRNA-depleted total RNA-seq libraries were prepared using the KAPA RiboErase kit for ribosomal RNA depletion together with the KAPA RNA HyperPrep Kit (Roche). Libraries were sequenced on an Illumina NovaSeq X Plus instrument in paired-end mode (2 × 151 bp, Phred+33 quality encoding) to a depth of approximately 35–40 million read pairs per sample (median ≈ 38 million). Library preparation and sequencing were performed by Genome Technologies at The Jackson Laboratory for Genomic Medicine (Farmington, CT, USA). Across libraries, a mean of ∼97% of reads aligned to the genome.

#### Read processing and quantification

Raw reads were processed with the nf-core/rnaseq pipeline (v3.23.0), (Ewels, Peltzer et al. 2020). Within the pipeline, adapter and quality trimming was performed with Trim Galore (Krueger 2026), reads were aligned to the human reference genome with STAR (Dobin, Davis et al. 2013), and transcript quantification was performed with Salmon (Patro, Duggal et al. 2017). The genome and annotation were the GENCODE v49 (Mudge, Carbonell-Sala et al. 2025) primary assembly. Salmon transcript-level estimates were summarized to gene level, and the merged gene-count matrix was used for all downstream analyses. The same GENCODE v49 annotation was used for gene-biotype and chromosome assignments throughout.

#### Sample-distance analysis

All downstream analyses were performed in R v4.3.1 (Team 2023) using DESeq2 v1.42.1 (Love, Huber et al. 2014). Gene counts were rounded to integers and imported into DESeq2. Low-count genes were removed (retaining genes with ≥ 10 counts in ≥ 3 samples), and counts were transformed with the variance-stabilizing transformation (VST, blind). Pairwise Euclidean distances were computed between samples on the transposed VST matrix, and the resulting distance matrix was clustered by hierarchical clustering with average linkage and visualized as a heatmap, annotated by lineage.

#### Differential expression and marker identification

Lineage markers were identified with DESeq2 using the negative-binomial Wald test. For each lineage we contrasted differentiated samples against the undifferentiated state (lineage vs Und), and to define a pluripotency signature we contrasted Und against all differentiated samples pooled (Und vs Differentiated). For each comparison, genes were filtered (≥ 10 counts in ≥ 3 samples) before fitting. Genes were considered differentially expressed at an adjusted *p* < 0.05 (Benjamini–Hochberg), |log2 fold change| > 1 and base mean > 75. Full and significant result tables of the top up-regulated genes were generated for every comparison (**Dataset S3**).

#### Protein-coding gene detection and CPM-threshold sensitivity

To quantify the breadth of the transcriptome captured by each lineage, counts were converted to counts per million (CPM) by dividing by per-sample library size. Protein-coding genes were defined by the protein_coding biotype in the GENCODE v49 primary-assembly annotation. Within each lineage, a gene was called expressed if it reached a given CPM threshold in at least two samples; the per-lineage expressed sets and their union across all lineages were then computed. To assess robustness of this definition, the analysis was repeated across a range of CPM thresholds (0.1, 0.25, 0.5, 1, 2, 5, 10, 25 and 50) and the number of expressed protein-coding genes was plotted as a function of threshold (log scale). A threshold of CPM > 1 was used for the primary expressed-gene definition.

#### Expressed genes call

Per-lineage expressed-gene calls were compiled into a supplementary table (**Dataset S5**) giving, for every annotated gene, its biotype, the mean CPM in each lineage, the number of samples in which it was expressed, and a per-lineage expressed/not-expressed call (CPM > 1 in ≥ 2 samples within the lineage), together with the number of lineages in which it was expressed. This resource allows the expression of any gene to be queried directly across lineages.

#### Inter-individual variability analysis

To highlight functional diversity across donors, we identified the genes most variable between individuals within each lineage (analysed where ≥ 3 individuals were available). To avoid sex-driven signal in a mixed-sex cohort, the analysis was restricted to autosomal genes (chrX, chrY and chrM excluded). For each lineage, a DESeq2 object was fit with a based on individual and two complementary metrics were combined: (i) a likelihood-ratio test (LRT) comparing *∼ Individual* against the reduced *∼ 1* model, and (ii) the median absolute deviation (MAD) and coefficient of variation of per-individual mean VST expression. Genes were ranked by the average of their MAD rank and LRT rank. Per-lineage variable-gene is supplementary material (**Dataset S4**).

#### Sample preparation and long-read sequencing

All genomic library preparation and sequencing described in this section were performed by Genome Technologies at The Jackson Laboratory for Genomic Medicine (Farmington, CT, USA).

#### Ultra-long Oxford Nanopore sequencing

High-molecular-weight (HMW) genomic DNA was extracted from 30 million cells using a phenol–chloroform protocol (Gong, Wong et al. 2019), followed by an additional purification with the Monarch HMW DNA Extraction Kit for Tissue (New England Biolabs, #T3060). Ultra-long libraries were prepared with the Ultra-Long DNA Sequencing Kit (SQK-ULK114, ONT) according to the manufacturer’s recommendations: 40 µg of DNA was incubated with 6 µL of fragmentation mix (FRA) at room temperature for 10 min and at 75 °C for 10 min, after which 5 µL of adapter (RAP-F) was added and the reaction incubated at room temperature for 60 min. Libraries were precipitated and resuspended in Elution Buffer (EB). Sequencing was performed on R10.4.1 flow cells (FLO-PRO114, ONT) on a PromethION (ONT) for 72–144 h, with multiple sequential loadings at approximately 24-h intervals or when pore activity dropped to 10–20%; a nuclease wash (Flow Cell Wash Kit, EXP-WSH004) was performed between successive loads.

#### PacBio HiFi sequencing

HMW DNA was extracted using the Monarch HMW DNA Extraction Kit for Cells and Blood (New England Biolabs, #T3050) according to the manufacturer’s instructions. Approximately 1– 2 million cells were first mildly lysed in Nuclei Prep Solution containing RNase A; Nuclei Lysis Solution containing Proteinase K was then added and the sample incubated at 56 °C for 10 min with agitation at 1,400 rpm. Genomic DNA was precipitated with isopropanol in the presence of glass beads on a rotating mixer at room temperature, washed twice with Wash Buffer, and eluted in Elution Buffer (PacBio) at 56 °C for 10 min; eluates were gently mixed with a wide-bore tip and held at 4 °C for 2–4 days to allow complete resuspension. Libraries were prepared with the SMRTbell prep kit 3.0 (PacBio): approximately 5 µg of HMW DNA was sheared on a Megaruptor 3 (Diagenode), cleaned with 1× SMRTbell cleanup beads, then end-repaired, A-tailed, and ligated to barcoded SMRTbell adapters, followed by cleanup, a nuclease treatment to remove fragments lacking adapters, and a further SMRTbell-bead purification. Libraries were size-selected on a PippinHT (Sage Science) to remove fragments < 10 kb and sequenced on a Revio (PacBio) with a 30-h movie.

#### Hi-C sequencing

Hi-C libraries were prepared with the Dovetail LinkPrep Kit (Cantata Bio) following the manufacturer’s instructions. One million cells were crosslinked with disuccinimidyl glutarate (DSG; Thermo Fisher Scientific) and formaldehyde (Sigma-Aldrich), after which in situ tagmentation and proximity ligation were used to capture spatially proximal DNA fragments. Crosslinks were then reversed and the DNA was purified. Libraries were generated with 12 cycles of PCR amplification, size-selected to 350–1,000 bp with AMPure XP beads (Beckman Coulter), and sequenced on an Illumina NovaSeq X Plus, generating 150 bp paired-end reads.

#### Haplotype-resolved genome assembly

For the eight GeDi lines and the KOLF2.1J reference, fully phased de novo assemblies were generated with hifiasm (v0.24.0) in Hi-C–integrated phasing mode, combining PacBio HiFi reads, ultra-long ONT reads, and Hi-C read pairs to resolve the two haplotypes and produce two haplotype assemblies (hap1, hap2) per line (**Figure S4**)

#### Assembly quality control

Assembly contiguity was summarized from the haplotype assemblies as the number and length distribution of contigs and the area under the Nx curve (auN). Gene completeness of the assemblies was assessed using compleasm (v0.2.7) using default parameters (Huang and Li 2023). BUSCO completeness is reported as the percentage of conserved single-copy orthologues from the primate lineage (primates_odb12) in OrthoDB v12 (**Figure 3A**). Compleasm estimates assembly completeness by identifying conserved single-copy orthologs and reports the proportion of complete (single and duplicated), fragmented, and missing genes. Errors in the assemblies were assessed using Flagger (v1.2.0). Reference representation was assessed by aligning each haplotype assembly to GRCh38 and computing the fraction of the genome covered and the contig spanning of each reference chromosome (**Figure 3C; Table S5**).

#### Variant calling

GRCh38 was used as the reference genome for all variant-calling analyses. Variant discovery was performed using complementary assembly-based, long-read, and short-read approaches.

Long-read and assembly-based variant calling (Figure S4): Structural variants (SVs) were identified from haplotype-resolved assemblies using PAV, SVIM-asm, and dipcall. For read-based SV calling, PacBio HiFi and Oxford Nanopore (ONT) reads were analyzed independently after alignment to GRCh38 using pbmm2 and minimap2, respectively. Sniffles2, longcallD, and DELLY were used to identify SVs from both PacBio HiFi and ONT data, whereas sawfish was applied exclusively to PacBio HiFi data. DELLY-CNV was used to identify copy-number variants (CNVs) from both HiFi and ONT data, while HiFiCNV was applied exclusively to HiFi data. SNVs and indels were identified from haplotype-resolved assemblies using PAV and dipcall and from PacBio HiFi and ONT reads using Clair3 and DeepVariant.

Short-read variant calling: Illumina short reads were aligned to GRCh38 using BWA-MEM and duplicated reads were marked and tagged using Picards MarkDuplicates step. SVs were identified using DELLY, SvABA, Manta, and SMOOVE, while SNVs were called using GATK. In addition, the Illumina DRAGEN pipeline was used for comprehensive variant calling, including SVs, CNVs, SNVs, and indels.

#### Variant merging and prioritization

VCF outputs from the 25 callers were normalized using bcftools, converting multiallelic records into biallelic records and left-aligning the position of indels within repeats. Bcftools was also used to remove variants which did not pass all caller filter criteria set during the calling process. Ensembl VEP (was used to annotate variants (using the --everything flag) up to 10 Mbp in size, associating them with genes up to 5000 bases away.

Ending coordinates were added to each record that reflect the size of the region affected by each variant, based on the length of the longer of the REF or ALT sequences. When the ALT sequence was not provided, caller-specific fields such as END, INSLEN, and SVLEN within the INFO fields were used to derive an ending coordinate. This ending coordinate was used for finding corroborating overlaps among different variants and overlaps between variants and repetitive regions that might give rise to artifacts during variant calling.

Variant calls were split into three groups: (1) SNVs; (2) non-SNV variants with affected regions less than or equal to 200 bases long (short variants), and variants with longer affected regions (long variants). SNV variants between different callsets were considered to corroborate one another if the chromosome, start position, and ALT sequence perfectly match. Short variant calls from different callers were considered to corroborate one another if the variant types were in the same compatibility set, if the reciprocal size match was 80% or greater, and if the variants were within 200 residues of one another. The four variant-type compatibility sets were {INS, DUP, CNV, BND}, {DEL, CNV, BND}, {INV, BND}, {SNV}. For example, SNVs can only match other SNVs, BND can match anything except an SNV, and DUP can match INS, CNV, or BND. Long variant calls from different callers were tabulated as corroborating one another if the variant types shared a compatibility set, and there was an 80% reciprocal overlap (overlap / max(len1, len2)) between the regions affected by each variant. BND variants were matched to the ends of other variants if the BND position was within three base pairs of either end of the other variant. The same matching rules were used when merging variants across samples.

Variants longer than 10 Mbp were removed, since several callers seem to make many false calls in this size range (data not shown). Calls of that size are more likely to be reliable if found using complementary technologies like G-banding or SNP arrays, at which point sequence data can be used to refine the ends of variants found by these latter methods.

Calls were filtered based on caller agreement. Only variants found by at least three callers, one of which had to be PAV were included. The variant structure determined by PAV was used as the exemplar variant for downstream annotation and reporting. Calls were removed if they overlapped by more than 75% with known UCSC problematic regions (hg38 version 2025-06-24, from https://hgdownload.soe.ucsc.edu/gbdb/hg38/problematic/comments.bb), or ENCODE exclusion regions (version 2020-05-05; from https://www.encodeproject.org/files/ENCFF356LFX/@@download/ENCFF356LFX.bed.gz). This process resulted in the final complete call set.

The complete call set was subsequently distilled into a high-priority list of variants most likely to affect cell line performance. We first removed variants whose VEP impact was not ‘HIGH’. High impact variants include those resulting in splicing variations, premature stop, lost start, frameshift, truncation or amplification, but does not include more moderate changes like synonymous and non-synonymous substitutions. Variants were also removed if they had a 1000Genomes allelic frequency greater than 1% (per VEP) and were not associated with pathogenicity by ClinVar (per VEP). Variants were also removed if they only affected pseudogenes, immunoglobulins, T-cell receptors, or individual ncRNAs. Variants affecting two or more distinct ncRNAs were kept in this priority set.

#### Structural variants in the PSG cluster and expression analysis

Structural variants in the *PSG* cluster were identified from the haplotype-resolved assemblies (PAV, v2.4.6). Two heterozygous intragenic deletions were analysed: a ≈63.9 kb deletion (chr19:43,197,628–43,261,491, hg38) that removes the promoter and 5′ exons of PSG4 and the body and 3′ exons of PSG9, joining the PSG9 promoter to the PSG4 3′ end (carriers GeDiX395, GeDiX927, GeDiY298), and a ≈18.9 kb deletion (chr19:42,909,001–42,927,865) that removes the promoter and 5′ exons of PSG6 and the 3′ exons of PSG7, joining the PSG7 promoter to the PSG6 3′ end (carrier GeDiX002).

Gene expression was quantified for the primitive-syncytium (PrS) condition (nine iPSC lines, three replicates each; DESeq2 variance-stabilising transformation, VST), with per-line values taken as the mean across replicates. To test whether the rearranged genes were specifically affected, each was compared to the structurally intact members of its own cluster: for each rearranged gene (PSG4, PSG9, PSG6, PSG7) we regressed its VST on a within-cluster reference — the per-line mean VST of the unrearranged cluster genes PSG1, PSG2, PSG3, PSG5 and PSG8 — using only non-carrier lines, and quantified each carrier’s deviation as the residual from this expectation.

##### Allelic DMR calling at FIRRE and the DXZ4 locus

For each female line, the active (Xa) and inactive (Xi) X chromosomes were assigned using the *XIST* regulatory DMR as anchor, defining Xi as the haplotype hypomethylated at this locus. Per-haplotype CpG methylation was summarized in 500-bp windows across each locus, and the Xa−Xi methylation difference was computed per window. Per-haplotype methylation was compared between the two alleles in fixed-size windows across each locus, and windows showing a consistent methylation difference between the active and inactive X were joined into DMRs and each DMR was classified as *concordant* (Xa > Xi, the same directionality as the *XIST* DMR) or *reversed* (Xi > Xa).

##### DXZ4 copy number from phased assemblies

Because the *DXZ4* macrosatellite lies within a 100 kb bridged gap in the hg38 reference, copy number was determined directly from the phased genome assemblies. For each haplotype, the assembled region corresponding to the DXZ4 locus was identified using the PAV alignment blocks flanking the reference gap as anchors, and a broad interval spanning these anchors was extracted from the contig. Within each extracted interval, the full extent of the DXZ4 array was then located de novo as a contiguous GC-rich block (∼0.62 GC) distinguishable from the surrounding lower-GC sequence. The length of the repeating unit was determined directly from each assembled array by measuring how far the sequence must be shifted along itself before it realigns — this shift corresponds to one repeat unit — and copy number was calculated as the total array length divided by that unit length. Calls were retained where the measured period fell within the expected *DXZ4* monomer range (∼3 kb) and the resulting array length and copy number fell within the reported population range (36–288 kb; 12–100 copies).

##### FIRRE local-repeat catalog and copy number

The *FIRRE* locus contains a family of local repeats (LRs) that are collapsed or incompletely represented in the short-read reference. For each X-bearing haplotype, the *FIRRE* locus was extracted from the phased assembly, and repeat families were characterized de novo with RepeatScout v1.0.6 (Price, Jones et al. 2005) and filtered to locus-restricted families by screening candidate consensus sequences against the full assembly (families dispersed genome-wide, i.e. transposable elements, were removed). Per-family copy number was counted by BLASTn (v2.14.1+) of each consensus against the haplotype, merging overlapping hits into distinct copies; apparent per-haplotype differences were confirmed by targeted realignment at reduced identity thresholds (80/70/60%). Families were reconciled across haplotypes by consensus length and matched to the reported catalog (Hacisuleyman, Shukla et al. 2016). The nuclear-retention repeat domain (RRD) is nested within a larger repeat and does not appear as an independent de novo family; it was recovered separately as the ∼150-bp unit recurring across the *FIRRE* exons and counted by BLASTn. Repeat instances were placed in reference coordinates by RepeatMasking (v4.1.9) the reference locus with the LR library and intersected with the allelic DMRs.

##### CTCF binding-site annotation at FIRRE

To assess whether the allelic DMRs coincide with CTCF binding, we used the ENCODE registry of candidate cis-regulatory elements (cCREs; ENCODE3, encodeCcreCombined, hg38) for the FIRRE locus. Elements carrying the CTCF-bound classification (CTCF-only elements and CTCF-bound enhancers and promoters) were retained as CTCF binding sites and intersected with the allelic DMRs. Because cCREs are aggregated across ENCODE biosamples, this annotation reflects CTCF binding observed in at least one cell type.

#### Haplotype-resolved methylation calling

To resolve methylation by haplotype, we employed the matched haplotype-resolved assemblies and their PAV-derived variant calls. Haplotype-resolved methylation was computed per chromosome as follows. For each chromosome, the PAV contig-alignment BEDs for haplotype 1 and haplotype 2 were intersected with the chromosome interval, and the corresponding assembly query regions (± 10 kb flank) were extracted from the hap1 and hap2 contigs to build a combined per-chromosome “mini-reference”. ONT reads aligning to that chromosome in the hg38 BAM were extracted, converted to FASTQ, and re-aligned to the mini-reference with minimap2 v2.31(Li 2018). Each read was assigned to haplotype 1 or haplotype 2 according to which haplotype the primary alignment mapped to; reads mapping to both haplotypes were flagged ambiguous and discarded.

Per-haplotype CpG methylation was called with modkit v0.5.1, producing bedMethyl in which the modified fraction is Nmod/Nvalid. Per-chromosome, per-haplotype bedMethyl files were concatenated genome-wide and coordinate-sorted. For visualization, the modified fraction was converted to bedGraph and then to bigWig with bedGraphToBigWig v2.10 (Kent, Zweig et al. 2010).

#### Global methylation across genomic features

For genome-wide and feature-level methylation, the per-chromosome, per-haplotype modkit bedMethyl files were first recombined into a single diploid CpG track per line. The feature classes analyzed were CpG islands, CpG shores, CpG shelves, promoters, gene bodies, and the LINE, SINE and LTR repeat classes, plus a genome-wide category. CpG islands were taken from the UCSC cpgIslandExt track (hg38) (Nassar, Barber et al. 2023); CpG shores and shelves were defined as the 2 kb regions immediately flanking islands (0–2 kb and 2–4 kb from the island edge, respectively). Promoters were defined as transcription start sites ± 2 kb (4 kb windows) and gene bodies as full gene-span intervals, both derived from the protein-coding genes of GENCODE v44 (Ensembl 110), (Mudge, Carbonell-Sala et al. 2025), one interval per gene (n = 20,033). LINE, SINE and LTR elements were extracted from the UCSC RepeatMasker track (rmsk) by repeat class (repClass field), (Nassar, Barber et al. 2023). To obtain a value for each feature class (e.g., SINE, LINE, CpG island) in each line, the combined CpG track was intersected with that class’s annotation, which returns every CpG overlapping any annotated element of the class. These CpGs were pooled across all elements of the class genome-wide. The genome-wide value used all CpGs in the track with no intersection. From the pooled CpG set for a line and feature class we then computed a coverage-weighted mean, in which each CpG contributes in proportion to its read depth (reported throughout), and, for reference, an unweighted mean, the arithmetic average of the per-CpG methylation percentages.

#### Comparison against reference methylomes (PCA)

To place the iPSC methylomes relative to public references, CpG methylation was binned into non-overlapping 10 kb genome-wide windows (mean methylation per window; windows with < 3 covered CpGs set to missing). The same binning was applied to publicly available whole-genome bisulfite methylomes: ENCODE blood references (CD14+ monocytes, CD3+ T cells, NK cells), (Consortium 2012) and Roadmap Epigenomics references spanning pluripotent (H1, H9, HUES64 ESC; iPS DF 6.9, iPS DF 19.11), somatic (liver, lung, spleen, IMR90) and hematopoietic (HSC CD34+) states (Roadmap Epigenomics, Kundaje et al. 2015). Windows covered in ≥ 80% of samples were retained; residual missing values were imputed with the per-window mean. The matrix was standardized (z-scored per window) and decomposed by principal component analysis (scikit-learn), (Pedregosa, Varoquaux et al. 2011).

#### Allelic methylation at imprinted DMRs

We assembled a curated set of 41 primary (germline) imprinted differentially methylated regions from published studies (Court, Tayama et al. 2014, Akbari, Garant et al. 2022, Jima, Skaar et al. 2022). DMR coordinates were confirmed against our own data by manual inspection of allelic methylation in IGV. For each line and DMR, mean methylation was extracted from the haplotype-1 and haplotype-2 bigWigs and used to compute (i) the absolute allelic methylation difference, Δ = |hap1 − hap2|, and (ii) the mean methylation of the two haplotypes, (hap1 + hap2)/2.

#### Allele-specific expression

Heterozygous SNVs for read phasing were obtained from the PAV VCFs. Using bcftools v1.12 we retained phased heterozygous SNVs on autosomes and chrX (chrY excluded as hemizygous in male lines), normalized against GRCh38, sorted and indexed. RNA-seq reads were merged across all six lineages (undifferentiated and five differentiation lineages) per line to maximize coverage, and allele-specific read counts were generated with phASER (Castel, Mohammadi et al. 2016) against the per-line heterozygous SNV set. Gene-level haplotypic counts were produced with phASER using a GENCODE v49 exon feature set in which exons were labeled by gene name and deduplicated.

Genes were classified following the binomial approach. Per gene, haplotypic counts were summed across exons; genes with ≥ 50 total reads were retained. The minor-allele fraction was defined as minor / (major + minor). A one-sided binomial test was applied against a monoallelic null (H0: minor-allele fraction = 0.1; H1: > 0.1), and p-values were corrected genome-wide by Benjamini–Hochberg FDR. Genes with FDR < 0.01 were called biallelic (rejecting the monoallelic null) and the remainder monoallelic (Topa, Benoit-Pilven et al. 2024).

---

**Supplementary Figure S1.**
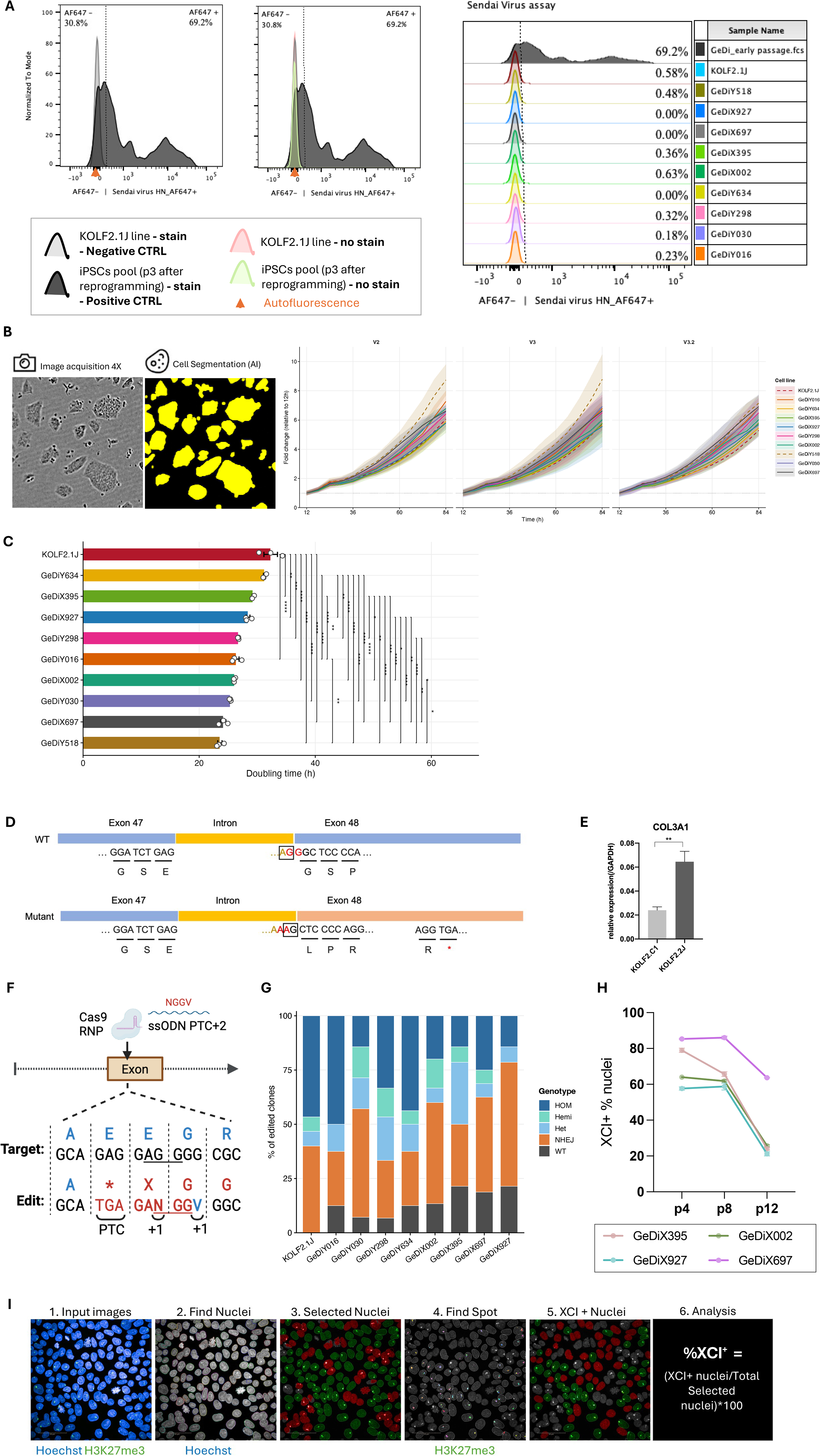
GeDi iPSC line growth and functional assay. **(A)** Sendai virus clearance assessed by flow cytometry using an anti-HN-AF647 antibody. Left panels show representative histograms for KOLF2.1J (negative control, unstained and stained) and the early-passage iPSC pool at passage 3 (positive control, unstained and stained). Right panel shows overlaid histograms for all nine GeDi lines and KOLF2.1J. Percentages indicate HN-AF647+ cells. All clonal lines show <1% positivity, confirming clearance of residual Sendai virus. **(B)** Representative phase-contrast images of GeDi colonies with corresponding AI-based cell segmentation masks used for IncuCyte confluency quantification (left). On the right, growth curves from three independent experiments showing fold change in confluency relative to the 12 h timepoint for all GeDi lines and KOLF2.1J. Shaded bands indicate 95% confidence intervals. **(C)** Doubling times for all GeDi lines and KOLF2.1J derived from IncuCyte growth curves across three independent experiments. Points represent individual experiments; bars show mean ± SD. Statistical significance was assessed by one-way ANOVA with Tukey post-hoc correction (*p < 0.05, **p < 0.01, ***p < 0.001). **(D)** Schematic of *COL3A1* mutation. GG at the splicing junction of exon 48 were mutated to AA, resulting in novel AG splicing site marked in rectangular. Only one allele showed this mutation in KOLF2.C1 (parental line) and its ARID2 mutant-corrected descendent KOL2.1J. **(E)** Expression of *COL3A1* in both KOLF2.C1 and KOLF2.2J derived ExMC, quantified by RT-qPCR. In KOLF2.2J cell line, this *COL3A1* mutation was corrected by CRISPR/Cas9. Error bar was shown as SD. Statistical test was done by t-test. * P_value<0.05, **P_value<0.01, ***P_Value<0.001. **(F)** Schematic of the PTC+2 CRISPR strategy used for GCM1 knock-out. A Cas9 RNP and ssODN donor were used to introduce a single-nucleotide edit converting a GGC (Gly) codon to a TGA stop codon with a +1 frameshift (PTC+1), generating a premature termination codon. The biallelic PTC+1 genotype is confirmed by Sanger sequencing (bottom). **(G)** Editing outcomes across all GeDi lines and KOLF2.1J following GCM1 PTC+1 knock-out. Stacked bar plots show the proportion of genotypes among 16 clones per line: homozygous (HOM), hemizygous (Hemi), heterozygous (Het), NHEJ, and wild-type (WT). Homozygous knock-out clones were recovered from all lines. **(H)** Longitudinal tracking of XCI maintenance in the four female GeDiX lines from passage 4 to passage 12. Each line shows the percentage of XCI+ nuclei (H3K27me3 focus-positive) at passages 4, 8, and 12. All four lines show progressive XCI erosion with increasing passage number. **(I)** Image analysis pipeline for automated XCI quantification. Representative images illustrate each step: (1) input image (Hoechst + H3K27me3 channels), (2) nucleus detection, (3) selected nuclei, (4) H3K27me3 spot detection, (5) XCI+ nuclei (nuclei containing at least one H3K27me3 focus), and (6) calculation of %XCI+ as (XCI+ nuclei / total selected nuclei) × 100.

**Supplementary Figure S2.**
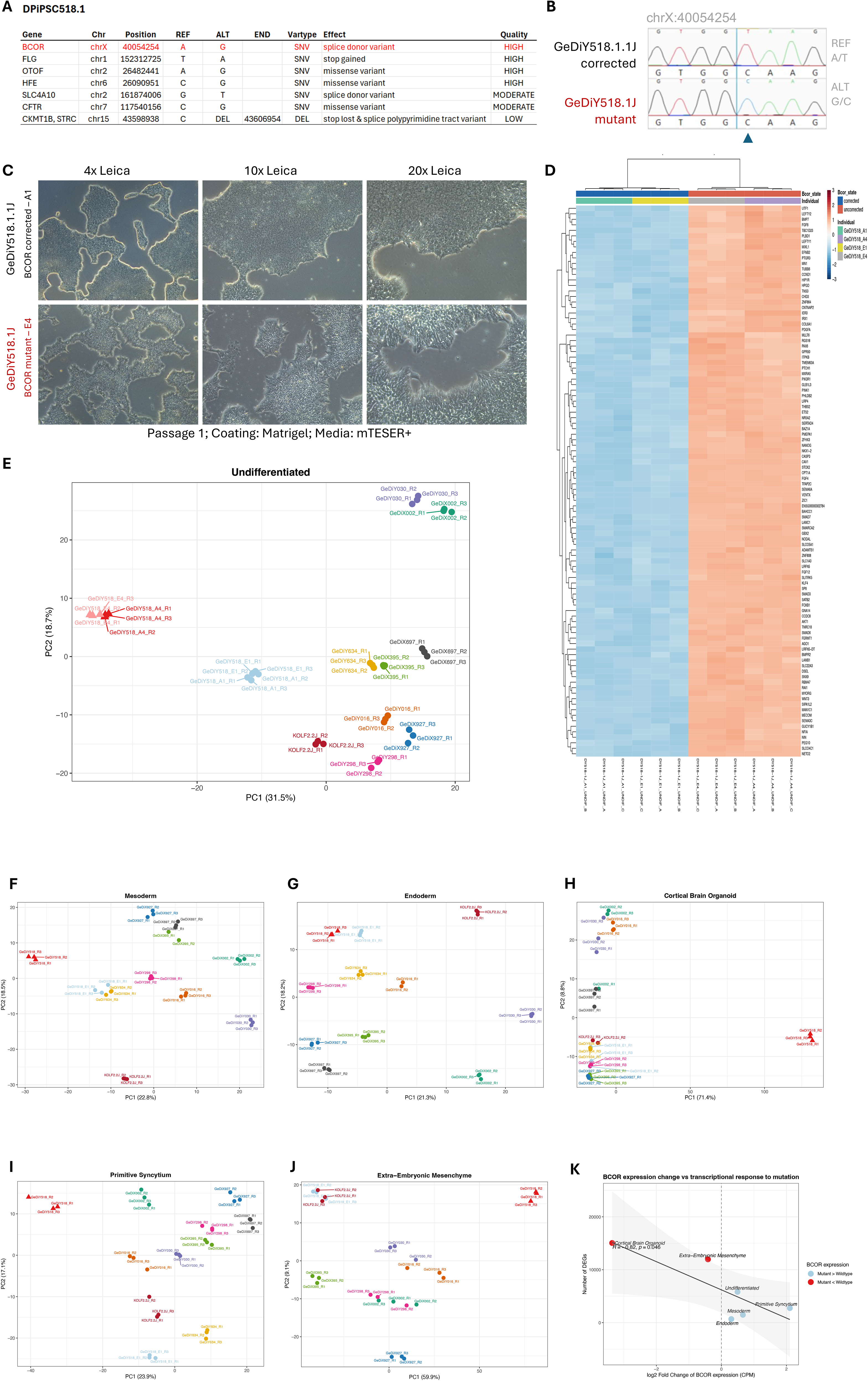
GeDiY518.1J BCOR loss-of-function variant. **(A)** Summary of high- and moderate-impact variants identified by Illumina short-read sequencing of GeDiY518.1J (DPIPSC518.1). The BCOR splice donor variant (chrX:40054254, A>G) is highlighted in red as the highest-priority variant. Additional variants in FLG, OTOF, HFE, SLC4A10, CFTR, and CKMT1B/STRC are listed with their chromosomal position, reference and alternate alleles, variant type, predicted effect, and quality classification. **(B)** Sanger sequencing confirmation of the BCOR splice donor variant and isogenic correction. Chromatograms show the chrX:40054254 locus in the isogenic corrected line GeDiY518.1.1J (REF A/T, top) and the BCOR mutant GeDiY518.1J (ALT G/C, bottom). The variant position is indicated by a blue arrowhead. **(C)** Brightfield images of GeDiY518.1.1J (BCOR corrected, top row) and GeDiY518.1J (BCOR mutant, bottom row) at passage 1 on Matrigel in mTeSR+ media, imaged at 4X, 10X, and 20X magnification (Leica). The BCOR mutant shows markedly altered colony morphology compared to the corrected line. **(D)** Heatmap of differentially expressed genes between GeDiY518.1J (BCOR mutant) and GeDiY518.1.1J (BCOR corrected) across replicate samples. Rows represent genes; columns represent individual samples. Colour scale indicates normalized expression (blue: low; red: high). Individual and Bcor state annotations are shown above. **(E)** PCA of RNA-seq data from undifferentiated GeDi samples across all lines (PC1: 31.5% variance; PC2: 18.7% variance). GeDiY518.1J (BCOR mutant, red ellipse) clusters separately from all other lines. The isogenic corrected line GeDiY518.1.1J (dashed ellipse) clusters with the broader GeDi panel, confirming that the transcriptomic divergence is driven by the BCOR variant rather than donor background. **(F–J)** Principal component analysis of differentiated lineages. Each panel shows PC1 versus PC2 for variance-stabilized transformed (VST) counts. Points are colored by individual iPSC line and shaped by BCOR state, with triangles indicating the BCOR-mutant line GeDiY518.1J and circles indicating all non-mutant lines, including the isogenic corrected control GeDiY518.1.1J. Each line is represented by three biological replicates. (F) Mesoderm. (G) Endoderm. (H) Cortical brain organoid. (I) Primitive syncytium. (J) Extra-embryonic mesenchyme. **(K)** Relationship between BCOR expression change and transcriptional response to mutation across lineages. Scatter plot showing the correlation between the log2 fold change of BCOR expression (mutation/no_mutation, calculated from median CPM values) on the x-axis and the number of differentially expressed genes (DEGs) identified in each lineage on the y-axis. Each point represents one differentiation lineage. Red points indicate lineages where the BCOR mutant shows lower expression than wildtype, blue points indicate lineages where the mutant shows higher BCOR expression than wildtype. The grey shaded area represents the 95% confidence interval of the linear regression. Pearson correlation coefficient and p-value are shown.

**Supplementary Figure S3.**
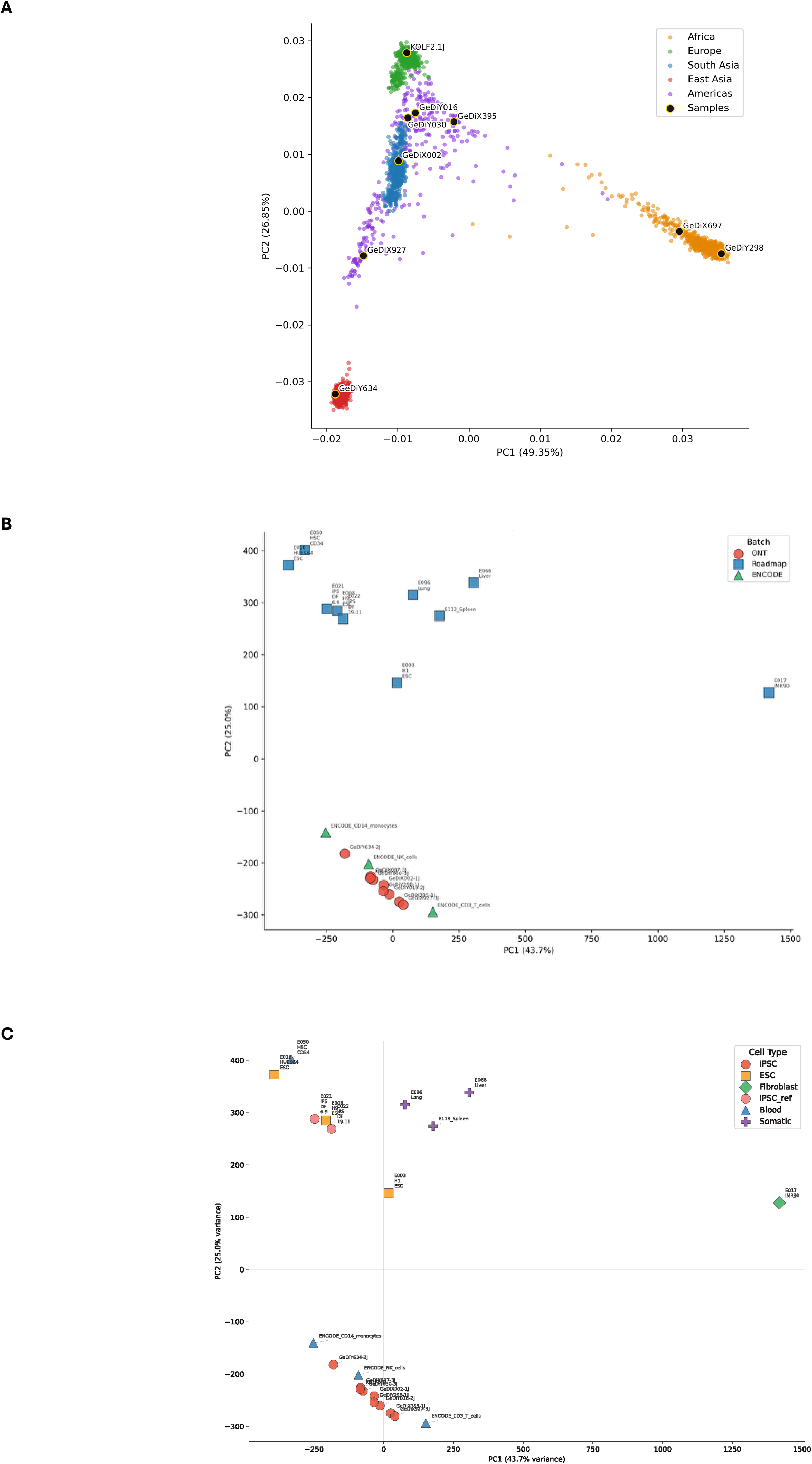
Complementary PCAs for ancestry and methylome analysis. **(A)** Projection of GeDi iPSC lines onto principal components 1 and 2 of the 1000 Genomes reference panel. SNP-array genotypes from the nine iPSC lines (black points with gold outlines) were projected onto principal components derived from the 1KG reference samples, colored by 1KG secondary label (AFR, orange; EUR, green; SAS, blue; EAS, red; AMR, purple). Axis labels give the percentage of variance captured by each component. PC2 separates the 1KG-EUR and 1KG-EAS reference samples but places the 1KG-SAS samples along the cline between them, with 1KG-AMR samples overlapping the same region; PC3 resolves these groups into distinct clusters and is shown in Figure 1D. GeDiY030 falls within the 1KG-AMR/SAS reference samples on this projection but is displaced from all reference groups on PC3, consistent with the absence of Middle Eastern populations from the 1KG panel. **(B)** Principal component analysis of the same genome-wide CpG methylation data (10 kb windows), shown as PC1 (43.7% of variance) versus PC2 (25.0% of variance), with symbols colored by data source (ONT, Roadmap, ENCODE). Along PC1 and PC2 the samples separate primarily by assay and data source — the long-read ONT methylomes (this study) resolve apart from the Illumina whole-genome bisulfite reference sets (Roadmap, ENCODE) — indicating that these components largely capture technical rather than biological variation. **(C)** The identical PC1-versus-PC2 projection as in (B), with symbols colored and shaped by cell type (GeDi iPSC, ESC, public iPSC, fibroblast, blood, somatic). Because PC2 is driven mainly by assay/platform, biological groupings (e.g. pluripotent versus blood) are not cleanly separated in this projection. This motivated the use of PC1 versus PC3 in the main figure, where the technical axis is excluded and samples resolve by biological state.

**Supplementary Figure S4.**
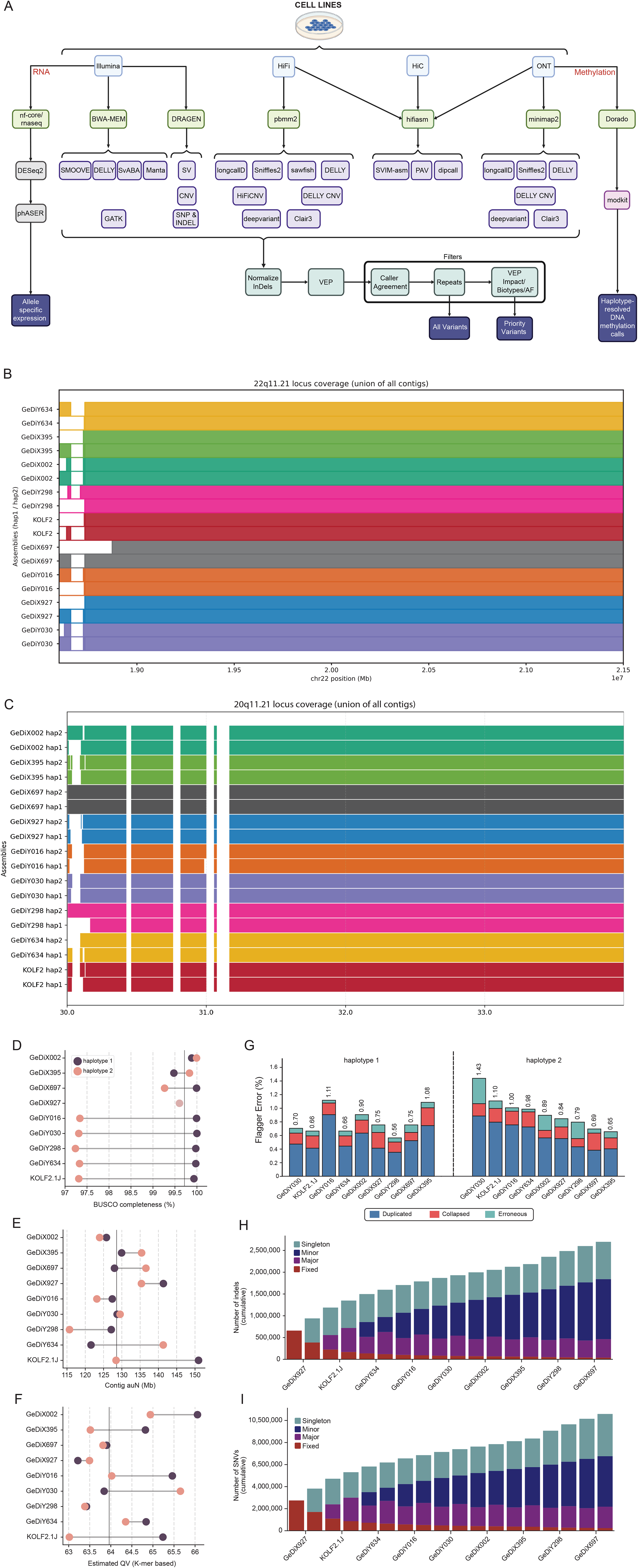
Haplotype-resolved genome assemblies — analysis pipeline and coverage at a structurally complex locus. **(A)** Schematic overview of the bioinformatic workflow for genome assembly, variant calling, gene expression analysis and DNA methylation analysis from iPSCs. Sequencing data included Illumina short-read RNA sequencing (RNA-seq) and whole-genome sequencing (WGS), Hi-C chromatin interaction data, PacBio high-fidelity (HiFi) long reads, and Oxford Nanopore Technology (ONT) long reads. RNA-seq data were processed with nf-core/rnaseq and DESeq2 for differential expression analysis and phASER for allele-specific expression. Short-read WGS data were aligned to GRCh38 using BWA-MEM and variants were identified using GATK, DRAGEN, SMOOVE, DELLY, SvABA, Manta. Haplotype-resolved genome assemblies were generated from HiFi, ONT and HiC data using hifiasm. Assembly-based variant calling was performed using PAV, SVIM-asm, and dipcall. Additionally, HiFi-reads and ONT-reads separately for variant detection using longcallD, Sniffles2, sawfish, DELLY, HiFiCNV, DELLY CNV, deepvariant, and Clair3. For DNA methylation profiling, ONT reads were basecalled using Dorado and analyzed with modkit to generate haplotype-resolved methylation calls. Final outputs include comprehensive non-redundant variant calls, priority variant calls, and haplotype-resolved DNA methylation profiles. **(B)** Assembly coverage across the 22q11.21 locus (chr22 position in Mb) for all GeDi lines and KOLF2 haplotypes (hap1 and hap2 shown separately per line). Each colored bar represents the union of contigs spanning this region for a given haplotype. White space indicates gaps regions in GRCh38. **(C)** Assembly coverage across the 20q11.21 locus (chr20 position in Mb) for all GeDi lines and KOLF2 haplotypes (hap1 and hap2 shown separately per line). Each colored bar represents the union of contigs spanning this region for a given haplotype. White space indicates gaps regions in GRCh38. (D) BUSCO completeness across all cell lines and corresponding haplotypes. Haplotypes containing chrY were designated as haplotype (hap2). (E) Assembly contiguity measured by area under the assembly length distribution curve (auN; Mb). Haplotypes containing chrY were designated as haplotype (hap2). (F) Assembly quality value (QV) scores across all cell lines and corresponding haplotypes. Haplotypes containing chrY were designated as haplotype (hap2). (G) Assembly quality assessment showing proportion of regions classified as duplicated, erroneous or collapsed in each assembly. Haplotypes containing chrY were designated as haplotype (hap2). (H) Cumulative non-redundant indels identified across all haplotypes relative to GRCh38. (I) Cumulative non-redundant SNVs identified across all haplotypes relative to GRCh38.

**Supplementary Figure S5.**
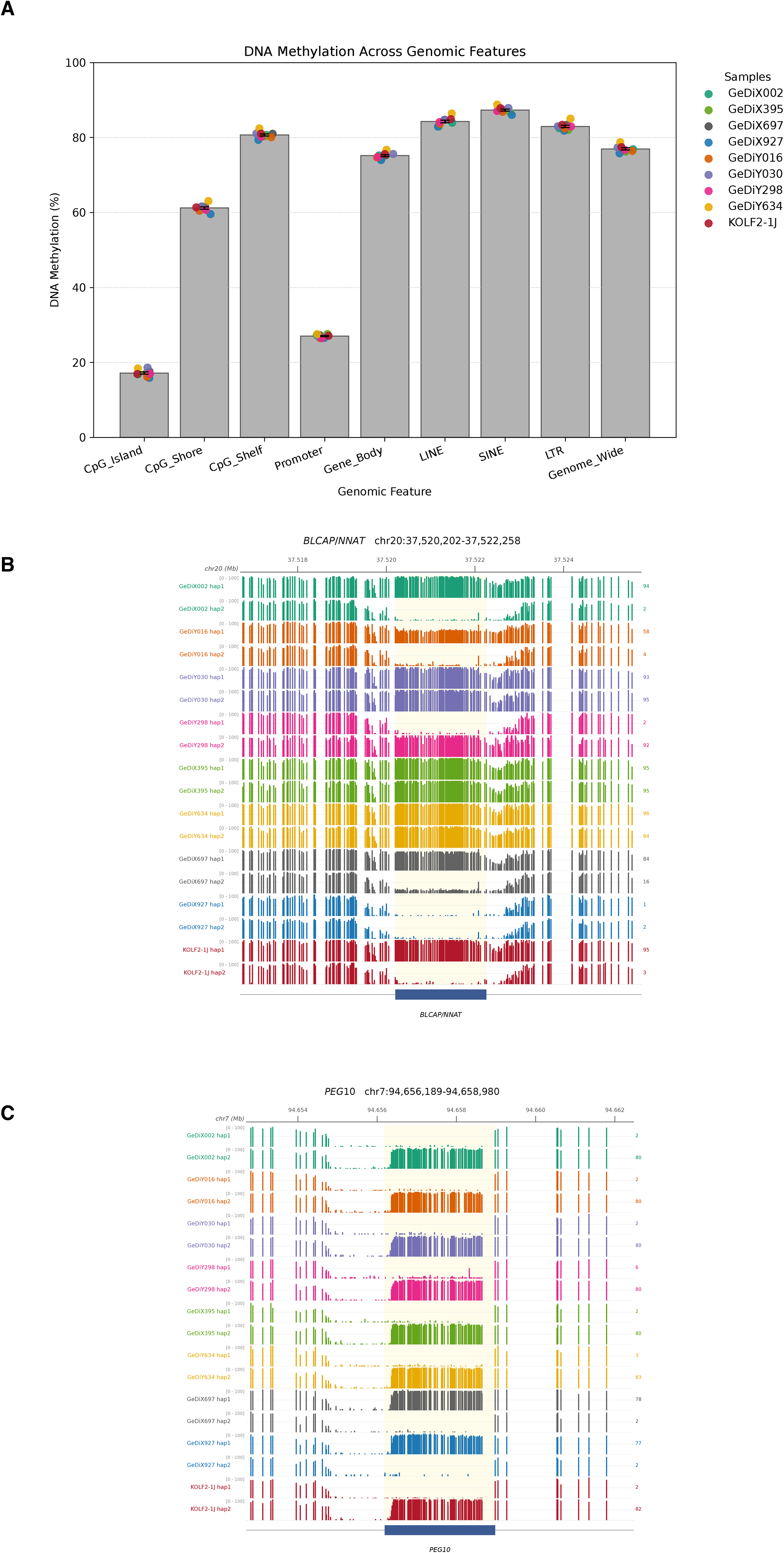
Genome-wide methylation integrity and haplotype-resolved methylation at representative imprinted DMRs. Panels show data for the nine iPSC lines (eight GeDi lines and the KOLF2-1J reference), except (D), which shows the four female lines. **(A)** DNA methylation across major genomic compartments. For each line, coverage-weighted mean CpG methylation was computed over CpG islands, CpG shores, CpG shelves, promoters, gene bodies, LINE, SINE and LTR elements, and genome-wide. Bars show the mean across lines, error bars the SEM, and colored dots the individual lines. **(B, C)** Haplotype-resolved CpG methylation at two imprinted DMRs. For each line, two tracks (hap1, hap2) display per-CpG methylation as vertical bars (bar height = methylation, 0–100%) across the locus; tracks are colored by line, the DMR interval is shaded, a genomic-coordinate ruler is shown at the top, and the value at the right of each track is the mean methylation across the DMR for that haplotype. **(B)** *BLCAP*/*NNAT* (chr20:37,520,202–37,522,258), a DMR with variable allelic methylation across lines. **(C)** *PEG10* (chr7:94,656,189–94,658,980), a DMR that retains the canonical imprinted configuration — one hypermethylated and one hypomethylated haplotype — across all lines.

## Supplementary Tables

- **Table S1.** SNP array
- **Table S2.** Emedgene variant calls — VUS and likely pathogenic/pathogenic variants
- **Table S3.** Ancestry proportions
- **Table S4.** Canonical lineage markers
- **Table S5.** Assembly metrics
- **Table S6.** All samples ASE results
- **Table S7.** All samples ChrX ASE results annotated
- **Table S8.** GeDi donor metadata
- **Table S9.** Variant count
- **Table S10.** Priority variant call

## Supplementary Datasets

- **Dataset S1.** Sample identity markers — transcriptomic (hg38)
- **Dataset S2.** Sample identity markers — genomic (hg38)
- **Dataset S3.** Upregulated genes across all comparisons
- **Dataset S4.** Allele-specific expression results across all samples
- **Dataset S5.** Variable gene expression across lineages
- **Dataset S6.** Expressed genes per lineage

